# Sweet revenge: AtSWEET12 in plant defense against bacterial pathogens by apoplastic sucrose limitation

**DOI:** 10.1101/2021.10.04.463061

**Authors:** Urooj Fatima, Muthappa Senthil-Kumar

## Abstract

Depriving bacterial pathogens of sugars is a potential plant defense strategy. The relevance of SUGARS WILL EVENTUALLY BE EXPORTED TRANSPORTERS (SWEETs) in plant susceptibility to pathogens has been established, but their role in plant defense remains unknown. We identified *Arabidopsis thaliana* SWEETs (AtSWEETs) involved in defense against nonhost and host *Pseudomonas syringae* pathogens through reverse genetic screening of *atsweet1–17* mutants. Double/triple mutant, complementation, and overexpression line analysis, and apoplastic sucrose estimation studies revealed that AtSWEET12 suppresses pathogen multiplication by limiting sucrose availability in the apoplast. Localization studies suggested that plant defense occurred via increased plasma membrane targeting of AtSWEET12 with concomitant AtSWEET11 protein reduction. Moreover, the heterooligomerization of AtSWEET11 and AtSWEET12 was involved in regulating sucrose transport. Our results highlight a PAMP-mediated defense strategy against foliar bacterial pathogens whereby plants control AtSWEET11-mediated sucrose efflux in the apoplast through AtSWEET12. We uncover a fascinating new mechanism of pathogen starvation as a broad-spectrum disease resistance mechanism in parallel with existing immune pathways.

**One sentence summary:** The transporter AtSWEET12 restricts bacterial pathogen multiplication by regulating sucrose availability to pathogens in the apoplast.

## INTRODUCTION

Most plant nutrients are sequestered inside the cell and are not easily accessible to bacterial pathogens^1,2^. Meanwhile, the apoplast is a nutrient niche in plant cells, where some nutrients are readily available. Many bacterial pathogens colonize the apoplast and utilize the sugars available therein. Studies in *Arabidopsis thaliana* and *Nicotiana benthamiana* have indicated that bacterial infection alters membrane permeability characteristics, leading to the release of sugars from the cytosol into the apoplast^3^. Plant defense mechanisms limiting sugar availability to bacterial pathogens have been ascribed to the regulation of sugar levels in the apoplast^3-8^. Thus far, studies related to plant defense mechanisms have mainly focused on antimicrobial compounds, reactive oxygen species production, the hypersensitive response, callose deposition, etc.^9-11^. However, the concept of sugar limitation to pathogens as a plant defense strategy has not been examined in detail.

Sugars are transported symplastically via plasmodesmata or apoplastically via sugar transporters located on the plasma membrane^6,12^. It is reasonable that plants might regulate sugar levels in the apoplast by controlling sugar transporters. A recent study in Arabidopsis suggested a plant defense strategy involving the control of sugar uptake in the apoplast by regulating sugar transporter protein 13 (STP13), which limits sugar availability to bacterial pathogens^13^. Further, a new class of sugar efflux transporters—SUGARS WILL EVENTUALLY BE EXPORTED TRANSPORTERS (SWEETs)—has been identified in several plant species^12,14,15^. In Arabidopsis, they comprise 17 members, which are divided into four clades^12,14-18^. The members of clades I, II, and IV are mainly hexose transporters, involved in the efflux of glucose, galactose, and fructose, respectively^7,15^. Clade III SWEET members preferentially transport sucrose^12^. AtSWEET11 and AtSWEET12 belong to this clade and are involved in sucrose efflux from the phloem parenchyma into the phloem apoplast, a critical step for subsequent phloem loading^6^.

Successful pathogens manipulate host plant sugar transporter machinery to redirect sugar efflux into the apoplast and establish their virulence^12,19^. Some SWEETs have been shown to be hijacked by bacterial pathogens for the release of sugar into the apoplast. For example, in rice, OsSWEET11 and OsSWEET14 are targeted by *Xanthomonas oryzae* pv. *oryzae* for releasing sugar into the apoplast^12,20,21^. Similarly, in Arabidopsis, the expression of different *AtSWEET* genes, including *AtSWEET11, AtSWEET12*, and *AtSWEET15*, is induced upon infection with fungi and hemibiotrophic bacterial pathogens^12^.

Contrarily, we speculate that as a part of the defense response, plants modulate the expression of *AtSWEET* genes to limit sugar release into the apoplast and thereby restrict sugar availability to bacterial pathogens, a likely explanation for “nonhost resistance.” Nonhost resistance is a durable type of plant disease resistance shown by an entire plant species to all isolates of a particular pathogen^22-24^. Hence, restricting sugar availability to nonhost pathogens is hypothesized to be an important plant defense strategy as a part of nonhost resistance^3,24^. In this study, we focused on 17 members of the AtSWEET family of sugar transporters to understand the broad defense mechanism that might act against nonhost pathogens, namely, *Pseudomonas syringae* pv. *phaseolicola* and *P. syringae* pv. *tabaci*, and the host pathogen *P. syringae* pv. *tomato* DC3000

## RESULTS

### Identification of the AtSWEET class of transporters involved in plant defense

The role of AtSWEETs belonging to four different clades has been elucidated in plant development and pathogenesis (Supplementary Table 1, Supplementary Fig. 1). To identify the AtSWEETs potentially involved in plant defense, we performed the reverse genetic screening of all 17 Arabidopsis *atsweet* mutants and tested their response to two nonhost pathogens, namely, *P. syringae* pv. *phaseolicola* (*Psp*) and *P. syringae* pv. *tabaci* (*Pta*). The bacterial multiplication assay indicated that compared to the corresponding wild-type plants, *atsweet12* and *atsweet15* mutants supported more *Psp* and *Pta* bacterial multiplication, while *atsweet17* mutants supported more *Psp* bacterial multiplication only (Fig. 1a, Supplementary Fig. 2). The *atsweet12* and *atsweet15* mutant plants demonstrated compromised nonhost resistance toward both the nonhost pathogens. However, the bacterial multiplication numbers for *Psp* and *Pta* in *atsweet11* mutant plants were significantly lower than those in wild-type plants (Fig. 1a, Supplementary Fig. 2).

**Fig. 1.**
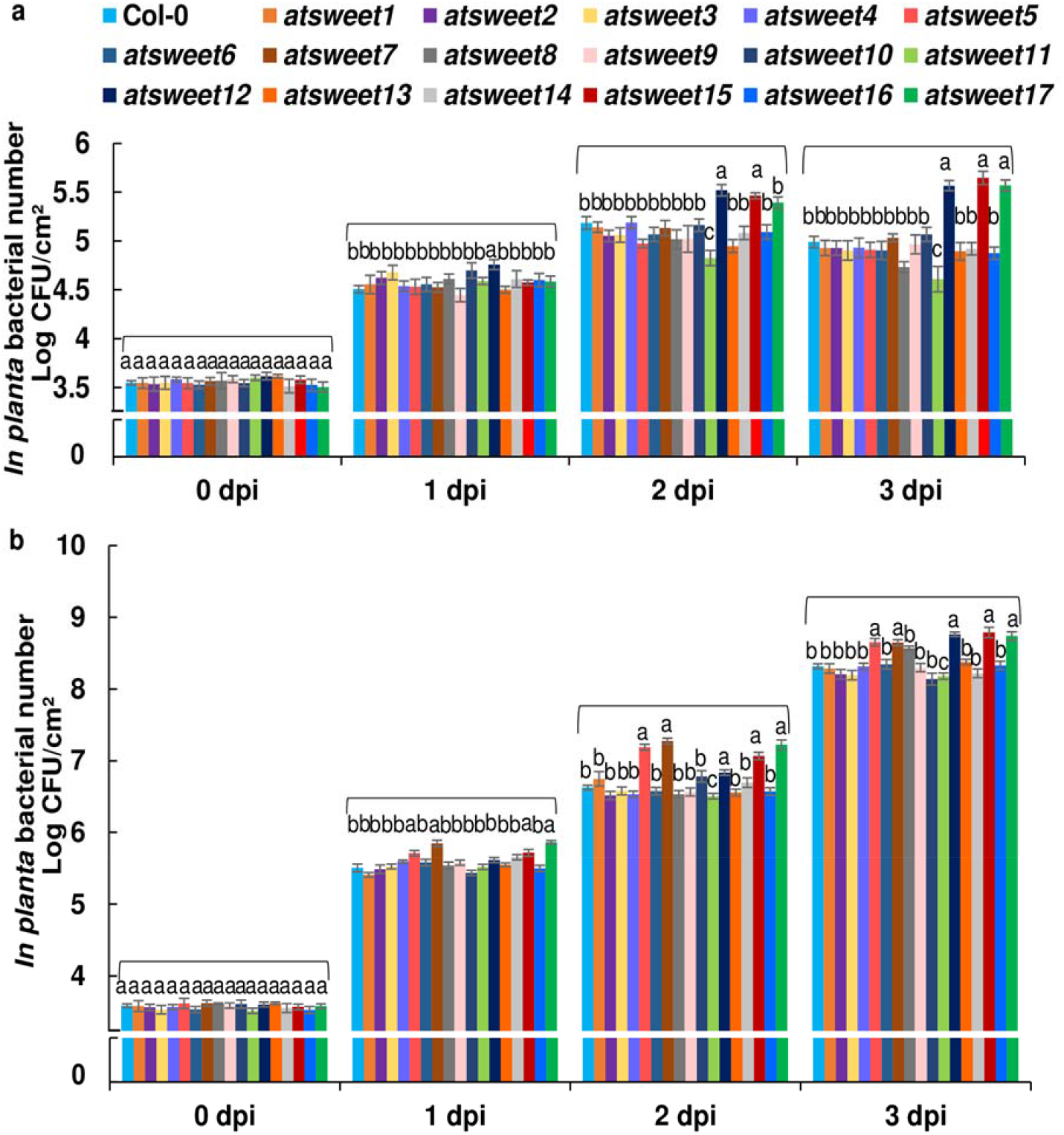
Mutant screening for the identification of AtSWEET transporters involved in plant defense. **a** and **b** The leaves of 32-day-old Arabidopsis wild-type (Col-0) and *atsweet1* to *atsweet17* mutant plants were inoculated with **(a)** the nonhost pathogen *Pseudomonas syringae* pv. *phaseolicola* (*Psp*) at 1 × 10^6^ CFU/mL and **(b)** the host pathogen *P. syringae* pv. *tomato* DC3000 (*Pst* DC3000) at 5 × 10^5^ CFU/mL. Bacterial multiplication assays were performed, and the bacterial populations were monitored by plating serial dilutions of leaf extracts at 0, 1, 2, and 3 days post inoculation (dpi). *In planta* bacterial number was expressed as log_10_ values. Significant differences (*P* < 0.05) after applying one-way ANOVA and Tukey’s correction are indicated by different letters. Data were obtained from the mean of six biological replicates and two technical replicates. Error bars show the standard error of the mean (SEM) (see Supplementary Dataset S1 for raw data and statistics). The experiment was repeated twice, and consistent results were observed.

We further examined whether AtSWEETs are involved in basal defense against a host pathogen, *P. syringae* pv. *tomato* DC3000 (*Pst* DC3000). The increased bacterial number compared to wild-type plants in the bacterial multiplication assay showed that *atsweet5, atsweet7, atsweet12*, and *atsweet15* mutant plants were hypersusceptible (Fig. 1b). Correspondingly, these mutant plants showed a high disease index and a phenotype of severe chlorosis followed by necrosis, unlike wild-type plants (Supplementary Figs. 3, 4). However, although *atsweet17* mutant plants showed a high bacterial number, no significant difference was observed in disease index or phenotype compared to wild-type plants (Fig. 1b, Supplementary Figs. 3, 4). Further, *atsweet11* mutant plants showed a slight resistance response, i.e., lower host bacterial multiplication, lower disease index, and less severe chlorosis than those of wild-type plants (Fig. 1b, Supplementary Figs. 3, 4). Meanwhile, a susceptible response was seen in *atsweet12* and *atsweet15* mutant plants against the host pathogen. The results obtained from both nonhost and host pathogen infection data together indicate that AtSWEET11, AtSWEET12, and AtSWEET15 could be potential candidates involved in plant defense.

To corroborate the findings from reverse genetic screening of *AtSWEET11, AtSWEET12*, and *AtSWEET15*, the transcript levels of these genes were analyzed upon nonhost and host pathogen infection by RT-qPCR at 16 and 24 hours post inoculation (hpi). *AtSWEET11* transcripts were induced at 16 h and 24 h after host pathogen *Pst* DC3000 infection, but no alteration in the transcript levels was observed after infection with the nonhost pathogens compared to the mock control (Supplementary Fig. 5a). Thus, the expression data along with reverse genetic screening of *atsweet11* mutants point to the involvement of AtSWEET11 in facilitating host pathogen infection. Moreover, the transcript levels of *AtSWEET12* and *AtSWEET15* were highly induced at 16 h after nonhost and host pathogen infection compared to the mock control. At 24 hpi, the transcript levels of *AtSWEET12* and *AtSWEET15* were still upregulated after host pathogen infection compared to the mock control. However, in case of inoculation with the nonhost pathogen at 24 h, the transcript expression of *AtSWEET12* was downregulated, but no significant difference was observed in the transcript expression of *AtSWEET15* compared to the mock control (Supplementary Fig. 5a). Thus, AtSWEET12 and AtSWEET15 participate in plant defense against both nonhost and host pathogens. To investigate whether the alteration in the expression of *AtSWEET11, AtSWEET12*, and *AtSWEET15* is type III secretion system (T3SS)-dependent, we used T3SS-deficient hrcC insertion mutants (*hrcC*^−^) of *Pst* DC3000 and *Pta* to inoculate the leaves of wild-type plants. The transcript levels of *AtSWEET11, AtSWEET12*, and *AtSWEET15* were found to be upregulated at 16 h after inoculation with *Pst* DC3000 *hrcC*^−^ T3SS secretion mutants compared to the mock control (Supplementary Fig. 5a). Similarly, after inoculation with *Pta hrcC*^−^, the transcript levels of *AtSWEET12* and *AtSWEET15* were upregulated compared to the mock control. Hence, the induction of the transcript expression of these genes did not depend on the effectors of T3SS. To then determine whether the transcript levels of *AtSWEET11, AtSWEET12*, and *AtSWEET15* are modulated by the plant during pathogen-associated molecular pattern (PAMP)-triggered immunity (PTI), we examined the expression of these genes in leaves inoculated individually with the PAMP elicitor FLG22 and crude PAMPs obtained from *Pst* DC3000, *Psp*, and *Pta*. Interestingly, we observed the downregulation of *AtSWEET11* and strong induction of *AtSWEET12* and *AtSWEET15* 3 hpi after treatment with FLG22 and crude PAMPs of *Pst* DC3000, *Psp*, and *Pta* (Supplementary Fig. 5b, c). It appears that after PAMP perception, the transcript expression of *AtSWEET11* is reduced, while that of *AtSWEET12* and *AtSWEET15* is induced by the plant. Thus, AtSWEET12 and AtSWEET15 are crucial in suppressing pathogen multiplication, whereas AtSWEET11 facilitates pathogen multiplication.

Furthermore, the expression of *AtSWEET12* and *AtSWEET15* was found to be downregulated in *atsweet11*, and that of *AtSWEET11* and *AtSWEET15* was found to be downregulated in *atsweet12* mutant plants compared to the wild type (Supplementary Fig. 6a, b). In comparison to the mocks, nonhost pathogen infection induced *AtSWEET12* and *AtSWEET15* expression in the *atsweet11* mutant and *AtSWEET11* and *AtSWEET15* expression in the *atsweet12* mutant (Supplementary Fig. 6c, d). Thus, in the absence of AtSWEET12, which is involved in plant defense, the expression of *AtSWEET11* is high, which might support increased pathogen multiplication.

### AtSWEET12 contributes to plant defense by suppressing pathogen multiplication

To clarify the role of *AtSWEET11, AtSWEET12*, and *AtSWEET15* in plant defense, we examined the response of the double mutants *atsweet11;12, atsweet11;15*, and *atsweet12;15* and the triple mutant *atsweet11;12;15* toward the nonhost pathogens *Psp* and *Pta* and the host pathogen *Pst* DC3000. The bacterial multiplication assay indicated that compared to the wild type, both the nonhost pathogens multiplied more in the *atsweet12;15* double mutant, relatively less in the *atsweet11;12;15* triple mutant, and very little in the *atsweet11;12* double mutant; no significant difference in bacterial number was observed in the *atsweet11;15* double mutant (Fig. 2a, Supplementary Fig. 7a). The nonhost pathogen *Psp*-infected *atsweet12;15* double mutant plants produced chlorotic disease symptoms at 3 days post inoculation (dpi) (Fig. 2c). Furthermore, the bacterial number of the host pathogen *Pst* DC3000 was also found to be highly increased in *atsweet12;15*, moderately decreased in *atsweet11;12;15*, and highly decreased in *atsweet11;12*; however, there was no significant difference in *atsweet11;15* compared to the wild type (Fig. 2b). *Pst* DC3000-infected *atsweet12;15* mutant plants showed a hypersusceptible response, indicated by enhanced chlorotic and necrotic disease symptoms and high disease index at 3 dpi compared to wild-type plants (Fig. 2c, d). In contrast, *Pst* DC3000-infected *atsweet11;12* mutant plants showed milder chlorotic symptoms and lower disease index that those in the wild-type plants (Fig. 2c, d). Interestingly, the *atsweet11;12* double mutant plants were highly resistant compared to single *atsweet11* mutant and wild-type plants. Thus, mutation in both *AtSWEET11* and *AtSWEET12* enhanced the reduction in bacterial multiplication. Contrary to this, mutation in both *AtSWEET12* and *AtSWEET15* boosted bacterial multiplication but without additive effects on disease susceptibility. Simultaneous mutation in *AtSWEET11* and *AtSWEET15*, however, did not show any significant difference in bacterial multiplication compared to wild-type plants. These results suggest that ATSWEET11 is epistatic over ATSWEET12 but not ATSWEET15, and AtSWEET11 and AtSWEET12 play a major part during pathogen infection.

**Fig. 2.**
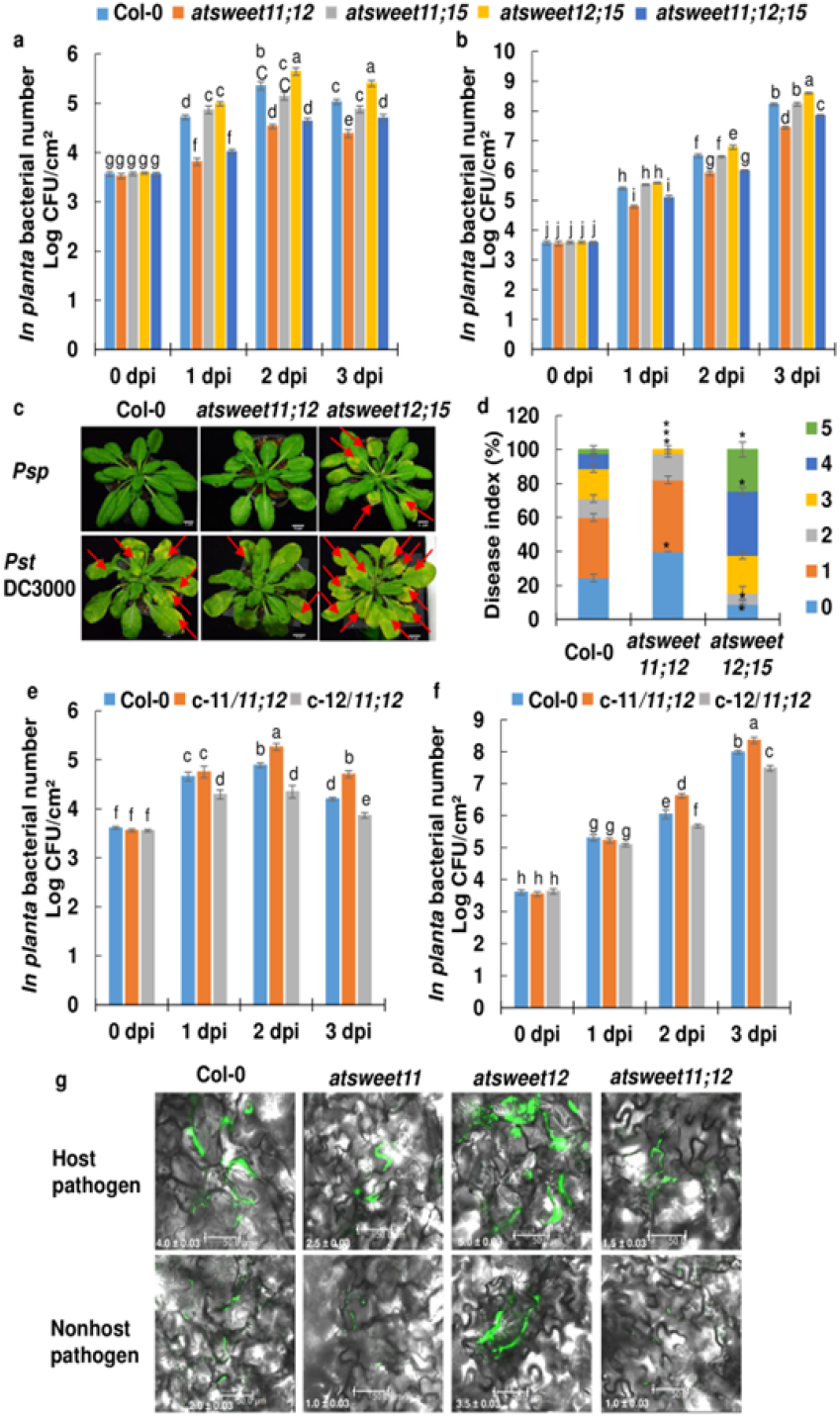
The sugar transporter AtSWEET12 is involved in plant defense. **a** and **b** The leaves of 32-day-old Arabidopsis wild-type (Col-0); *atsweet11;12, atsweet11;15*, and *atsweet12;15* double mutants; and *atsweet11;12;15* triple mutants were inoculated with **(a)** the nonhost pathogen *Psp* at 1 × 10^6^ CFU/mL and **(b)** the host pathogen *Pst* DC3000 at 5 × 10^5^ CFU/mL. Bacterial populations were monitored by plating serial dilutions of leaf extracts at 0, 1, 2, and 3 days post inoculation (dpi). **c** Phenotypes of plants inoculated with the *Psp* and *Pst* DC3000. Water-inoculated plants (mock treatment) were used for comparison with pathogen-inoculated plants. Photographs were taken at 3 dpi. **d** Disease scoring was done for host pathogen-infected plants at 3 dpi. Scores were given on the basis of symptom development as follows, score 0: no symptoms, score 1: leaf margins showing chlorosis, score 2: midrib region of leaf showing chlorosis, score 3: two-thirds of leaf area showing chlorosis, score 4: full leaf showing chlorosis, score 5: leaf showing necrosis. Asterisks indicate a significant difference from the wild-type (Student’s *t-*test; \**P* < 0.01). Data were obtained from the mean of three biological replicates, and error bars show the SEM. **e** and **f** Thirty-two-day-old Arabidopsis wild-type (Col-0) and complementation lines c-11/11;12 and c-12/11;12 transformed with p-AtSWEET11:AtSWEET11 and pAtSWEET12:AtSWEET12, respectively, in the *atsweet11;12* double mutant background were syringe inoculated with **(e)** *Psp* at 1 × 10^6^ CFU/mL and **(f)** *Pst* DC3000 at 5 × 10^5^ CFU/mL. Bacterial populations were monitored by plating serial dilutions of the leaf extract at 0, 1, 2, and 3 dpi. For a, b, e and f *in planta* bacterial number was expressed as log_10_ values. **g** The *in planta* bacterial populations of green fluorescent protein (GFP)-labeled host pathogen *Pst* DC3000 and nonhost pathogen *P. syringae* pv. *tabaci* (*Pta*) were determined in the wild-type (Col-0), *atsweet11, atsweet12*, and *atsweet11;12* mutant plants. The leaves of 32-day-old Arabidopsis wild-type, *atsweet11, atsweet12*, and *atsweet11;12* mutant plants were inoculated with the *Pst* DC3000 expressing *GFPuv* at 5 × 10^5^ CFU/mL and *Pta* expressing *GFPuv* at 3 × 10^5^ CFU/mL. The population of fluorescent bacteria was monitored at 2 dpi by observing the leaf using a Leica TCS SP8 confocal microscope (excitation at 488 nm and emission between 500 and 600 nm). The images were taken using a 63X objective and merged together by Leica microsystems LAS AF confocal software. The GFP fluorescence signals were quantified by ImageJ software (http://imagej.nih.gov/ij/). The intensity values were calculated and presented at the bottom of the image. The intensity was depicted as log_10_ values. For **a, b, e**, and **f**, significant differences (*P* < 0.05) after applying two-way ANOVA and Tukey’s correction are indicated by different letters. Data were obtained from the mean of six biological replicates and two technical replicates. Error bars show the SEM. The experiment was repeated thrice, and consistent results were observed.

The absence of AtSWEET11 and AtSWEET12 together led to a resistance response toward the host pathogen. Therefore, we assessed the response of the complementation lines transformed with either *p-AtSWEET11:AtSWEET11* or *pAtSWEET12:AtSWEET12* in *atsweet11;12* double mutant plants toward the nonhost pathogens *Psp* and *Pta* and the host pathogen *Pst* DC3000. The *in planta* bacterial number in the complementation line of *p-AtSWEET11:AtSWEET11* in the *atsweet11;12* double mutant background showed increased *Psp, Pta*, and *Pst* DC3000 bacterial multiplication than that in the wild-type plants (Fig. 2e, f, Supplementary Fig. 7b). The complementation line of *pAtSWEET12:AtSWEET12* in the *atsweet11;12* double mutant showed less *Psp, Pta*, and *Pst* DC3000 bacterial multiplication than that of the wild type (Fig. 2e, f, Supplementary Fig. 7b). It is evident that the susceptible response of the complementation lines of *p-AtSWEET11:AtSWEET11* in the *atsweet11;12* double mutant was similar to that of *atsweet12* single mutants, and the resistance response of the complementation line of *pAtSWEET12:AtSWEET12* in the *atsweet11;12* double mutant was similar to that of the *atsweet11* single mutant. Thus, in the absence of *AtSWEET12, AtSWEET11* appears to participate in enhancing the bacterial number, which leads to plant susceptibility. These results point to AtSWEET12 as one of the important players involved in plant defense against pathogen infection.

*In planta* green fluorescent protein (GFP)-labeled *Pst* DC3000 and *Pta* populations were monitored at 2 dpi. We found that the apoplast of *atsweet12* mutant plants supported high levels of *Pst* DC3000 and *Pta* populations compared to the wild type (Fig. 2g). However, the apoplast of *atsweet11* mutant plants supported almost similar levels of *Pst* DC3000 populations but fewer *Pta* populations than those of wild-type plants (Fig. 2g). Moreover, in the apoplast of *atsweet11;12* double mutant plants, *Pst* DC3000 and *Pta* populations were much lower than those in wild-type plants (Fig. 2g, Supplementary Fig. 8). Overall, our data from the microscopic studies corresponded with the results obtained from the bacterial multiplication assay in mutant and wild-type plants, indicating that the presence of AtSWEET12 is crucial in suppressing bacterial populations in the apoplast.

To determine whether mutation in *AtSWEET11* and *AtSWEET12* impacts the *in planta* multiplication of T3SS *hrcC*^−^ mutants of *Pst* DC3000, *Psp*, and *Pta*, as well as avirulent strains of *Pst* DC3000 (*AvrRpt2, AvrRps4*, and *AvrRpm1*), bacterial multiplication assays were performed in mutant and wild-type plants. Similar to the response observed for *Pst* DC3000, *Psp*, and *Pta*, the *in planta* bacterial number of *hrcC*^−^ mutants of these pathogens was lower in *atsweet11* and *atsweet11;12* mutants and higher in the *atsweet12* mutant compared to the wild type (Supplementary Fig. 9). These results indicate that the enhanced susceptibility of *hrcC*^−^ strains in *atsweet12* mutants is not dependent on effectors. Moreover, in the case of avirulent strains, no significant difference was observed in the bacterial multiplication number of *Pst* DC3000 (*AvrRpt2, AvrRps4*, and *AvrRpm1*) in *atsweet11* and *atsweet12* mutants compared to the wild type (Supplementary Fig. 10). This result suggests that the absence of *AtSWEET11* or *AtSWEET12* does not impact the *R* gene-mediated defense response against avirulent pathogens carrying the corresponding *avr* genes.

### Sucrose limitation in the mutant apoplast affects bacterial multiplication

To understand the relation between the bacterial multiplication pattern and sucrose availability, apoplastic sugar levels were estimated. Consistent with the involvement of AtSWEET11 and AtSWEET12 in sucrose transport, the relative and absolute apoplastic sucrose levels in the leaves of *atsweet11* and *atsweet11;12* mutant plants were half of those in wild-type plants (Fig. 3a, Supplementary Fig. 11). However, in the *atsweet12* mutant plants, the apoplastic sucrose levels were indistinguishable from those of the wild type (Fig. 3a, Supplementary Fig. 11). We further confirmed the difference in the sucrose levels in the apoplast obtained from mutant and wild-type plants by measuring the specific luminescence using lux-based sucrose biosensors^25^. We found that the specific luminescence intensity was lower in the apoplast from *atsweet11* and *atsweet11;12* mutant plants than that from wild-type and *atsweet12* mutant plants (Fig. 3b). Thus, the reduced accumulation of sucrose in the apoplast of *atsweet11* and *atsweet11;12* mutant plants concurred with the decreased bacterial multiplication in these mutant plants. In addition, the mutation in *AtSWEET11* reduced apoplastic sucrose levels, but the mutation in *AtSWEET12* alone showed no alteration in sucrose levels, indicating that AtSWEET11 might be actively participating in sucrose transport, unlike AtSWEET12.

**Fig. 3.**
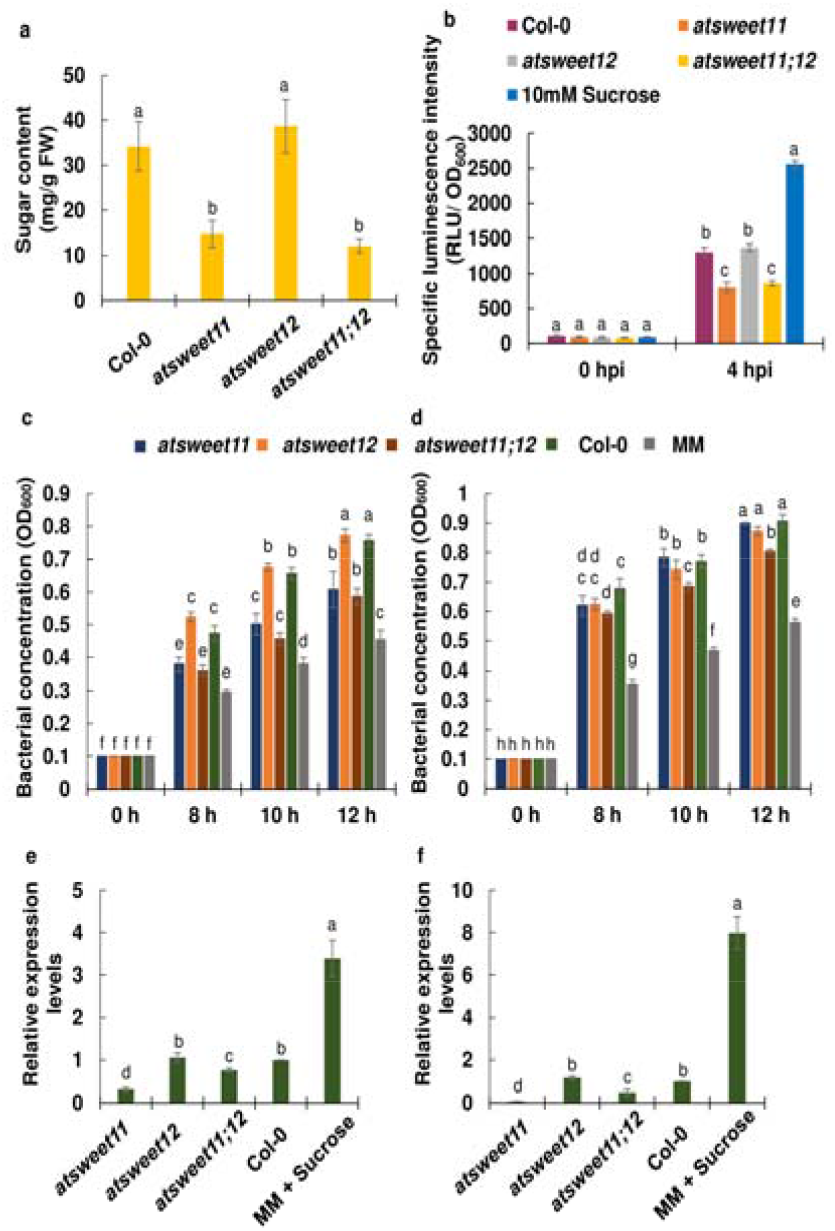
Apoplastic sucrose levels in mutant and wild-type plants impact the *in vitro* bacterial multiplication and sucrose utilization by bacteria. **a** The absolute level of apoplastic sucrose was measured by a sugar quantification kit. **b** The estimation of specific luminescence intensities of sucrose biosensors from *R. leguminosarum* grown in the apoplast obtained from *atsweet11, atsweet12, atsweet11;12* and wild-type (Col-0) plants. Bacteria were inoculated in the apoplast extracts such that the OD_600_ at 0 hour time point was maintained at 0.1 (OD_600_ = 0.1). Bacterial cultures were incubated at 28 °C, and relative luminescence (expressed as RLU) and OD_600_ was measured at 0 and 4 hpi. The specific luminescence intensity was calculated as RLU/ OD_600_. The 10mM sucrose was used as a positive control. **c** and **d** *In vitro* quantification of **(c)** *Psp* and **(d)** *Pst* DC3000 multiplication in apoplast extract of *atsweet11, atsweet12, atsweet11;12* and wild-type (Col-0). The apoplast was extracted from 32-day-old mutant and wild-type plants by the vacuum-infiltration and centrifugation method. Bacteria were inoculated in minimal medium M9 (MM) supplemented with 5 % apoplast extracts such that the OD_600_ at 0 hour time point was maintained at 0.1 (OD_600_ = 0.1). Bacterial cultures were incubated at 28 °C, and OD_600_ was measured at 0, 8, 10, and 12 hpi. **(e)** and **(f)** The transcript levels of the *scrY* gene (gene ID: *Psp*, PSPPH5187; *Pst* DC3000; PSPTO0890) were studied in **(e)** *Psp* and **(f)** *Pst* DC3000 grown *in vitro* in MM supplemented with apoplast extract of *atsweet11, atsweet12, atsweet11;12*, and wild-type (Col-0), and sucrose (50 mM). The bacteria were inoculated in MM supplemented with 5 % apoplast extracts and sucrose at 50 mM concentration (OD_600_ = 0.1 at 0 hpi). The bacterial culture was incubated at 28 °C. The bacterial cells were harvested at 10 hpi and bacterial RNA was isolated. The relative transcript levels were measured by RT-qPCR. The expression values were normalized against the reference gene, *16S rRNA* (*Psp*: PSPPH_0689; *Pst* DC3000: PSPTOr01), and the relative expression levels (RQ) were obtained over bacteria grown in Col-0 apoplast. Bars represent the transcript expression pattern of genes. In **a, b, e** and **f**, significant difference (*P* < 0.05) after applying one-way ANOVA and Tukey’s correction are indicated by different letters. For **a, e** and **f** data were obtained from the mean of three biological replicates (n = 3), and error bars show the SEM. The experiment was repeated twice, and consistent results were observed. For **b**, data were obtained from the mean of seven biological replicates (n = 7) and two technical replicates, and error bars show the SEM. The experiment was repeated thrice, and consistent results were observed. In **c** and **d** significant differences (*P* < 0.05) after applying two-way ANOVA and Tukey’s correction are indicated by different letters. The mean of five biological replicates (n = 5) was obtained, and error bars show the SEM. The experiment was performed thrice, with consistent results.

Minimal medium M9 (MM) supplemented with apoplastic fluids from the leaves of *atsweet11* and *atsweet11;12* mutant plants supported less *Psp* and *Pta* multiplication compared to the wild type (Fig. 3c, Supplementary Fig. 12). No significant difference was observed for *Psp* and *Pta* multiplication in apoplastic fluids from the *atsweet12* mutant compared to wild-type plants (Fig. 3c, Supplementary Fig. 12). Moreover, for *Pst* DC3000, *in vitro* bacterial multiplication in apoplastic fluid was found to be similar in the leaves of *atsweet11* and *atsweet12* mutants and the wild type. However, the apoplastic fluids from *atsweet11;12* leaves supported less *Pst* DC3000 multiplication compared to the wild type (Fig. 3d). These results indicate that less sucrose availability in the apoplast impedes *in vitro* bacterial multiplication. Moreover, the *in vitro* bacterial multiplication of *Psp, Pta*, and *Pst* DC3000 in MM supplemented individually with sucrose, glucose, and fructose indicated that sucrose and glucose were the preferred energy sources for these pathogens (Supplementary Fig. 13). Overall, the *in vitro* bacterial quantification studies clearly suggested that the mutation in *AtSWEET11* reduced sucrose levels and *in vitro* bacterial multiplication in the apoplast. However, the mutation in *AtSWEET12* did not impact the sucrose levels or *in vitro* bacterial multiplication in the apoplast. Hence, during plant defense, AtSWEET12 is involved in controlling the AtSWEET11-mediated sucrose efflux in the apoplast to restrict pathogen multiplication.

The bacterial gene *scrY* (*Psp*: PSPPH5187; *Pst* DC3000: PSPTO0890) encodes sucrose porin, which is involved in sucrose uptake inside bacterial cells. We performed a comparative analysis of the sucrose utilization ability of the bacterial pathogens in the apoplast obtained from mutant and wild-type plants by studying the transcript level of *scrY. scrY* expression in *Psp* and *Pst* DC3000 grown in apoplast extracts from *atsweet11* and *atsweet11;12* mutants was significantly lower than that in wild-type plants (Fig. 3e, f). However, no significant difference was observed in the expression level of this gene in *Psp* and *Pst* DC3000 grown in apoplast extracts from the *atsweet12* mutant compared to the wild type. Moreover, the expression of *scrY* in *Psp* and *Pst* DC3000 grown in MM supplemented with sucrose was significantly higher than that in MM supplemented with the apoplast extract from wild-type plants (Fig. 3e, f). This shows the correlation between sucrose availability in the apoplast and the sucrose utilization ability of the bacterial pathogens owing to the expression of genes involved in sucrose uptake. The low expression level of bacterial sucrose porin gene and low sucrose utilization ability of bacteria in the mutant apoplast indicate the low sucrose availability in the apoplast from the mutant which led to less bacterial multiplication.

### Plant defense deprives nonhost pathogens by restricting sucrose supply in the apoplast

To understand the impact of pathogen infection on the dynamics of apoplastic sucrose levels, we studied the sucrose levels of apoplastic fluids obtained at 24 hpi from mock-treated and pathogen-infected leaves of *atsweet11* and *atsweet12* mutants and wild-type plants. As expected, the apoplastic sucrose levels were significantly reduced in wild-type plants after nonhost pathogen infection compared to the mock treatment (Fig. 4a). Besides, in *atsweet11* mutant plants, apoplastic sucrose levels remained unaltered after nonhost pathogen infection (Fig. 4a). However, in *atsweet12* mutant plants, the apoplastic sucrose levels increased significantly after nonhost pathogen infection compared to the mock treatment (Fig. 4a, b). This suggests that plant defense against nonhost pathogens involves the function of AtSWEET12 in limiting sucrose availability in the apoplast to suppress pathogen multiplication. Further, in case of host pathogen infection, the apoplastic sucrose levels were significantly reduced in the infected leaves of the *atsweet11* and *atsweet12* mutants and wild-type plants compared to the mock treatment (Fig. 4a). Based on this result, we speculate that the reduction in apoplastic sucrose levels in any genotype after host pathogen infection might be the result of sugar acquisition by the host pathogen for nutrition^12^.

**Fig. 4.**
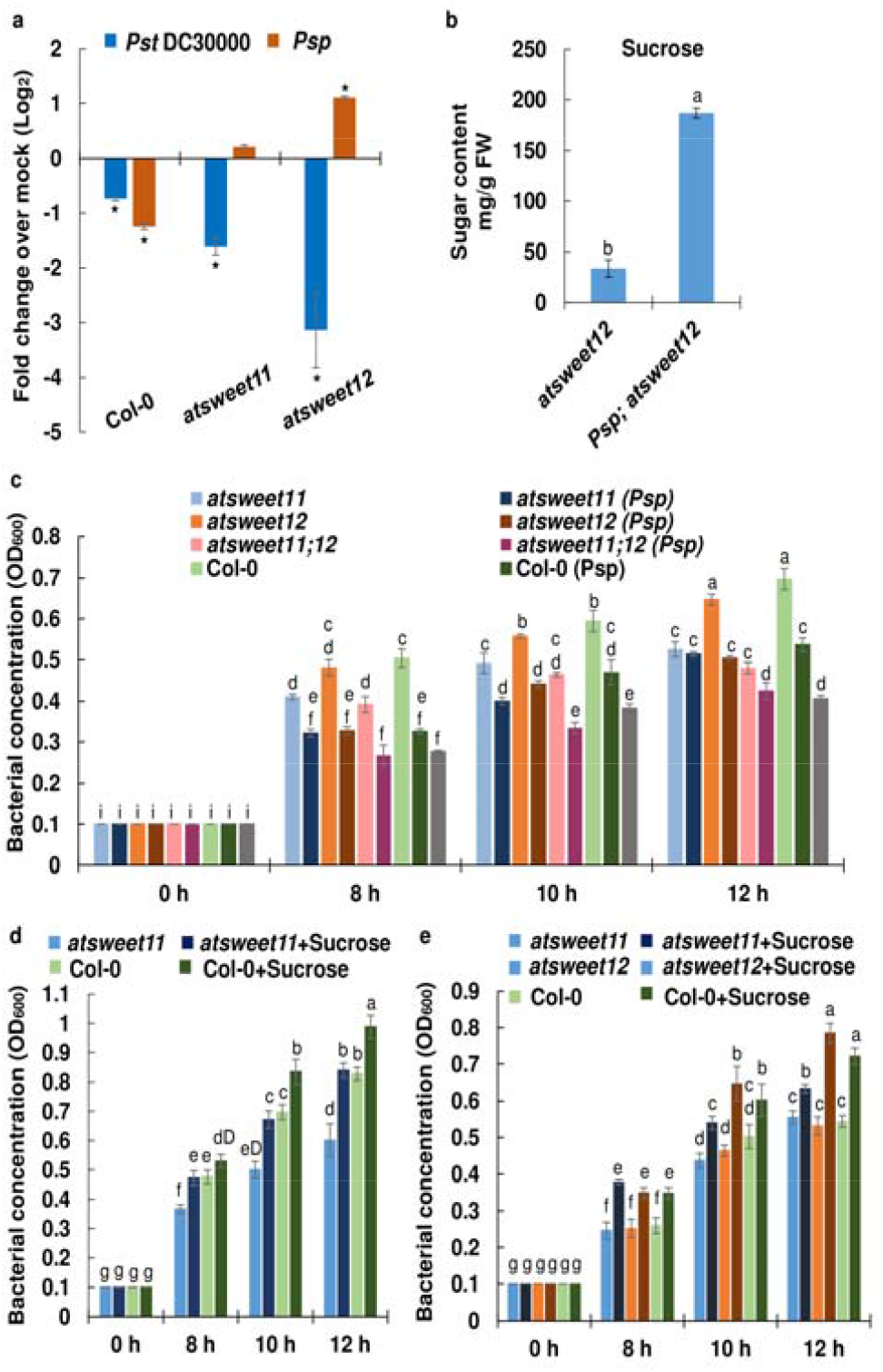
Sucrose is the limiting factor responsible for restricting the nonhost pathogen multiplication in the apoplast. **a** The apoplastic sucrose levels after host and nonhost pathogen infection in wild-type, *atsweet11* and *atsweet12* mutant plants over their respective mock-treated plants are presented. The 32-d-old Arabidopsis *atsweet11, atsweet12* mutant and wild-type plants were inoculated with sterile water as mock, host pathogen *Pst* DC3000 at 5 × 10^5^ CFU/mL, and nonhost pathogen, *Psp* at 1 × 10^6^ CFU/mL. Samples were harvested at 24 hpi. The apoplastic fluids from wild-type and mutant plants were obtained by vacuum infiltration and centrifugation method. The sucrose levels was estimated by using gas chromatography-mass spectrometry (GC-MS). The relative abundance of sugars were estimated by using ribitol as an internal standard. **b** The absolute sucrose content was measured in the apoplast obtained from *atsweet12* mutant plants inoculated with sterile water as mock and *Psp* at 1 × 10^6^ CFU/mL at 24 hpi by sugar estimation kit. **a, b** Asterisks indicate a significance difference from respective mock-treated plants (student’s *t* test; \**P* < 0.05). Data were obtained from the mean of total three biological replicates (n=3) and error bars show ± standard error of mean. The experiment was repeated twice and consistent results were observed. **c** *In vitro* quantification of *Psp* multiplication was done in the apoplast extract obtained at 8 h after infection with *Psp* and sterile water in *atsweet11, atsweet12, atsweet11;12*, and wild-type (Col-0) plants. The apoplastic extracts were isolated by the vacuum-infiltration and centrifugation method. Bacteria were inoculated in MM supplemented with 5 % apoplast extracts (OD_600_ = 0.1 at 0 hpi). The bacterial culture was incubated at 28 °C, and OD_600_ was measured at 0, 8, 10 and 12 hpi. **d** *In vitro* quantification of *Psp* multiplication in the apoplast extract of *atsweet11* and wild-type (Col-0) plants after 1 % sucrose addition in the apoplast. The apoplast was extracted from 32-day-old mutant and wild-type (Col-0) plants by vacuum-infiltration and centrifugation. **e** *In vitro* quantification of *Psp* multiplication after sucrose addition in the apoplast extract obtained 8 hpi with *Psp* and sterile water (used as the mock treatment) in *atsweet11, atsweet12*, and wild-type (Col-0) plants. The apoplast was extracted from *Psp* and mock-treated mutant and wild-type (Col-0) plants by vacuum-infiltration and centrifugation. *Psp* was inoculated in MM supplemented with 5 % apoplast extract and 1 % sucrose (OD_600_ = 0.1 at 0 hpi). Bacterial cultures were incubated at 28 °C, and OD_600_ was measured at 0, 8, 10, and 12 hpi. For **c, d** and **e** significant differences (*P* < 0.05) after applying two-way ANOVA and Tukey’s correction are indicated by different letters. Data were obtained from the mean of five biological replicates (n = 5), and error bars show the SEM. The experiment was repeated thrice, and consistent results were observed.

Next, we wanted to confirm whether plants restrict the sucrose availability in the apoplast as a defense response against nonhost bacterial pathogens. *In vitro* bacterial quantification was performed for *Psp* and *Pst* DC3000 individually in MM supplemented with apoplastic fluid extracted from wild-type and mutant (*atsweet11, atsweet12, atsweet11*;12) leaves infected with *Psp* and *Pst* DC3000, respectively. Apoplastic fluids from nonhost pathogen-infected leaves of mutant and wild-type plants were found to support less multiplication of the nonhost pathogen than the apoplastic fluids from the respective mock-treated leaves (Fig. 4c). However, in the case of host pathogens, no significant difference was observed (Supplementary Fig. 14). These results suggest that plants limit sugar levels in the apoplast as a defense response against nonhost pathogens, eventually preventing *in vitro* multiplication due to sugar deficiency in the apoplastic fluids.

To determine whether relieving the sucrose limitation in the apoplast extract from mutants can recover *in vitro* bacterial multiplication, we quantified bacteria after externally adding sucrose to the apoplast extracts. The sucrose supplementation in the apoplast extracts of *atsweet11* mutant leaves supported more *Psp* multiplication compared to the apoplast extracts of *atsweet11* mutant leaves without external sucrose addition. Interestingly, *Psp* multiplication in the apoplast of the *atsweet11* mutant with external sucrose was found to be similar to that observed in the apoplast from wild-type plants (Fig. 4d). These results indicate that the addition of sucrose in the apoplast of the *atsweet11* mutant relieved the sucrose limitation caused by the absence of AtSWEET11 and restored *in vitro* bacterial multiplication similar to the level observed in the apoplast from wild-type plants.

We also verified whether sucrose limitation in the apoplast from nonhost pathogen-infected plants was responsible for reduced *in vitro* bacterial multiplication and could be recovered by sucrose addition. We found that the external sucrose supplementation in apoplast fluids from nonhost pathogen-infected leaves of *atsweet11* and *atsweet12* mutant and wild-type plants supported more *Psp* multiplication *in vitro* compared to the respective mutant and wild-type apoplast fluids without sucrose (Fig. 4e). These results further suggest that plants limit sucrose availability to nonhost pathogens in the apoplast to restrict nonhost pathogen multiplication, and external sucrose supplementation in the apoplast restores *in vitro* bacterial multiplication.

### *In planta* sucrose addition restored bacterial multiplication in mutants to wild-type levels

We next explored whether sucrose is the limiting factor for reduced *in planta* bacterial multiplication in *atsweet11* and *atsweet11;12* mutant plants. After *in planta* addition of 0.2% sucrose, both *atsweet11* and *atsweet11;12*, supported more *Psp* multiplication compared to mutant plants without sucrose treatment (Fig. 5a). Interestingly, after sucrose addition, *Psp* multiplication in these mutant plants was significantly higher than in wild-type plants without sucrose treatment at 3 dpi (Fig. 5a). Similarly, after the addition of 1% sucrose in the *atsweet11* and *atsweet11;12* mutant plants, *Psp* multiplication was significantly higher, and chlorotic disease symptoms were observed, indicating higher plant susceptibility compared to mutant and wild-type plants without exogenous sucrose treatment (Fig. 5a, b). Despite *Psp* being a nonhost pathogen, the addition of sucrose in wild-type plants enhanced *Psp* multiplication compared to wild-type plants without sucrose treatment. These results indicate that the resistant phenotype of the *atsweet11* and *atsweet11;12* mutant and wild-type plants was reverted to the susceptible phenotype after *in planta* sucrose addition, and that sucrose is the limiting factor for nonhost pathogen multiplication in the apoplast of these mutant and wild-type plants. In case of *atsweet12* mutants, only at a higher concentration of sucrose, *Psp* multiplication was increased compared to the *atsweet12* mutant without sucrose treatment (Fig. 5a). This indicates that due to the absence of AtSWEET12, the AtSWEET11-mediated sucrose efflux is not controlled, and enough sucrose was available to the nonhost pathogen in the apoplast of *atsweet12* mutants; therefore, exogenous sucrose addition only at higher concentrations increased the multiplication of the nonhost pathogen.

**Fig. 5.**
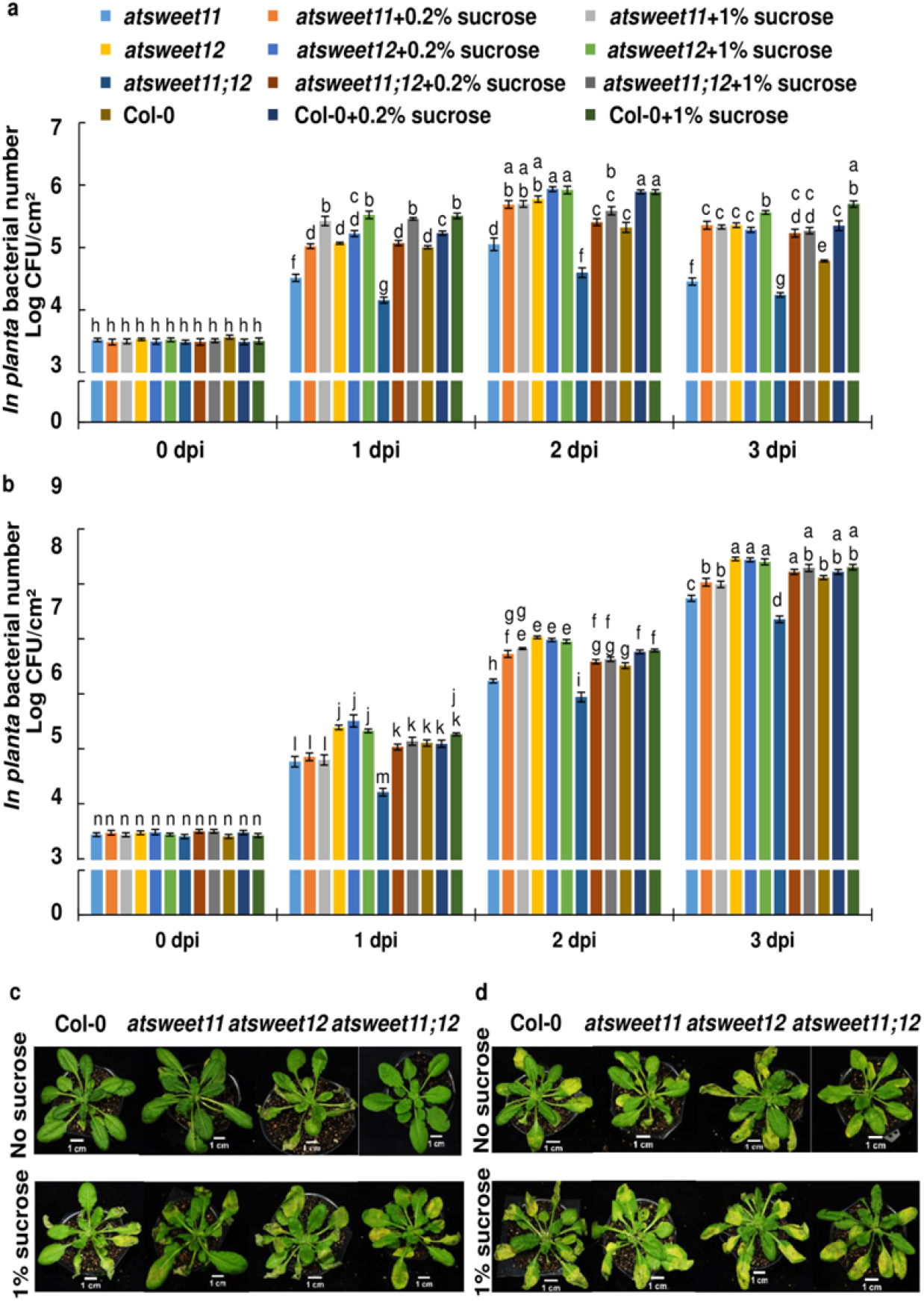
Pathogen multiplication in mutants is reverted to wild-type levels upon *in planta* sucrose addition. **a** *In planta* bacterial multiplication was estimated in 32-day-old Arabidopsis *atsweet11, atsweet12*, and *atsweet11;12* mutants and wild-type plants inoculated with the nonhost pathogen *Psp* alone, co-inoculated with *Psp* and 0.2 % sucrose, and co-inoculated with *Psp* and 1 % sucrose. **b** *In planta* bacterial multiplication was estimated in 32-day-old *atsweet11, atsweet12*, and *atsweet11;12* mutants and wild-type plants inoculated with the host pathogen *Pst* DC3000 alone, co-inoculated with *Pst* DC3000 and 0.2 % sucrose, and co-inoculated with *Pst* DC3000 and 1 % sucrose. For **(a)** and **(b)**, the bacterial population was monitored by plating serial dilutions of leaf extracts at 0, 1, 2, and 3 dpi. Significant differences (*P* < 0.05) after applying two-way ANOVA and Tukey’s correction are indicated by different letters. Data were obtained from the mean of six biological replicates and two technical replicates. Error bars show the standard error of mean (SEM). The *in planta* bacterial number was expressed as log10 values. **c** Phenotypes of mutant and wild-type plants inoculated with the nonhost pathogen *Psp* alone and co-inoculated with *Psp* and 1 % sucrose. **d** Phenotypes of mutant and wild-type plants inoculated with the host pathogen *Pst* DC3000 alone and co-inoculated with *Pst* DC3000 and 1 % sucrose. For **(c)** and **(d)**, photographs were taken at 3 dpi. The experiment was repeated twice, and consistent results were observed.

In the case of the host pathogen *Pst* DC3000, exogenous sucrose addition both at 0.2% and 1% concentration in *atsweet11;12* supported more *Pst* DC3000 multiplication compared to mutant and wild-type plants without sucrose treatment. Correspondingly, the *atsweet11;12* mutants after sucrose addition at a higher concentration showed severe chlorosis compared to mutant and wild-type plants without sucrose treatment (Fig. 5d). In the case of *atsweet11* mutants with exogenous sucrose addition, a significant increase in *Pst* DC3000 multiplication was observed at 2 dpi, and more chlorosis was seen compared to the *atsweet11* mutant and wild-type plants without sucrose treatment (Fig. 5b, d). Together, these results clearly indicate that sucrose was the limiting factor for host pathogen multiplication in *atsweet11* and *atsweet11;12* mutant plants, and exogenous sucrose addition alleviated the sucrose limitation in the apoplast, thereby leading to increased pathogen multiplication. Moreover, in *atsweet12* mutant and wild-type plants, exogenous sucrose addition at high or low concentrations did not show significant differences in *Pst* DC3000 multiplication compared to *atsweet12* mutant and wild-type plants without sucrose treatment, respectively (Fig. 5b). Accordingly, we surmise that in *atsweet12* and wild-type plants, sucrose availability might be adequate for a host pathogen to multiply in the apoplast; therefore, exogenous addition of sucrose in these plants did not cause any difference in pathogen multiplication.

### Plasma membrane targeting of AtSWEET12 with a concomitant reduction in AtSWEET11 led to plant defense

Since our study indicated the differential role of AtSWEET11 and AtSWEET12 in plant defense, we next assessed the localization pattern of AtSWEET11 and AtSWEET12 protein in Arabidopsis plants carrying *AtSWEET11:AtSWEET11-GUS* and *AtSWEET12:AtSWEET12-GUS* transgenes after nonhost and host pathogen infection. Histochemical localization by the GUS reporter assay indicated that AtSWEET11 transporters were well expressed in the vascular tissue, including the major and minor veins, of mock-treated *AtSWEET11:AtSWEET11-GUS* transgenic plants (Fig. 6a). After *Psp, Pta*, and *Pst* DC3000 infection, the expression of AtSWEET11 protein was reduced, as indicated by the reduction in GUS staining in the major and minor veins at 36 hpi and 48 hpi (Fig. 6a, Supplementary Figs. 15a, c, 16a). In contrast, the expression of AtSWEET12 protein was barely detected in the major and minor veins of mock-treated plants carrying *AtSWEET12:AtSWEET12-GUS* transgenes (Fig. 6b). However, *AtSWEET12* transcripts were well expressed under normal conditions (Supplementary Fig. 5c). The difference in transcript and protein level might be due to posttranslational regulation of transporters^26^. Moreover, after bacterial infection, irrespective of host and nonhost pathogen, AtSWEET12 protein expression was induced, as indicated by GUS staining in the major and minor veins at 36 hpi and 48 hpi (Fig. 6b, Supplementary Fig. 15b, d, 16b). Thus, protein plant defense actively involves the presence of AtSWEET12 with concomitant suppression of AtSWEET11 to restrict bacterial multiplication.

**Fig. 6.**
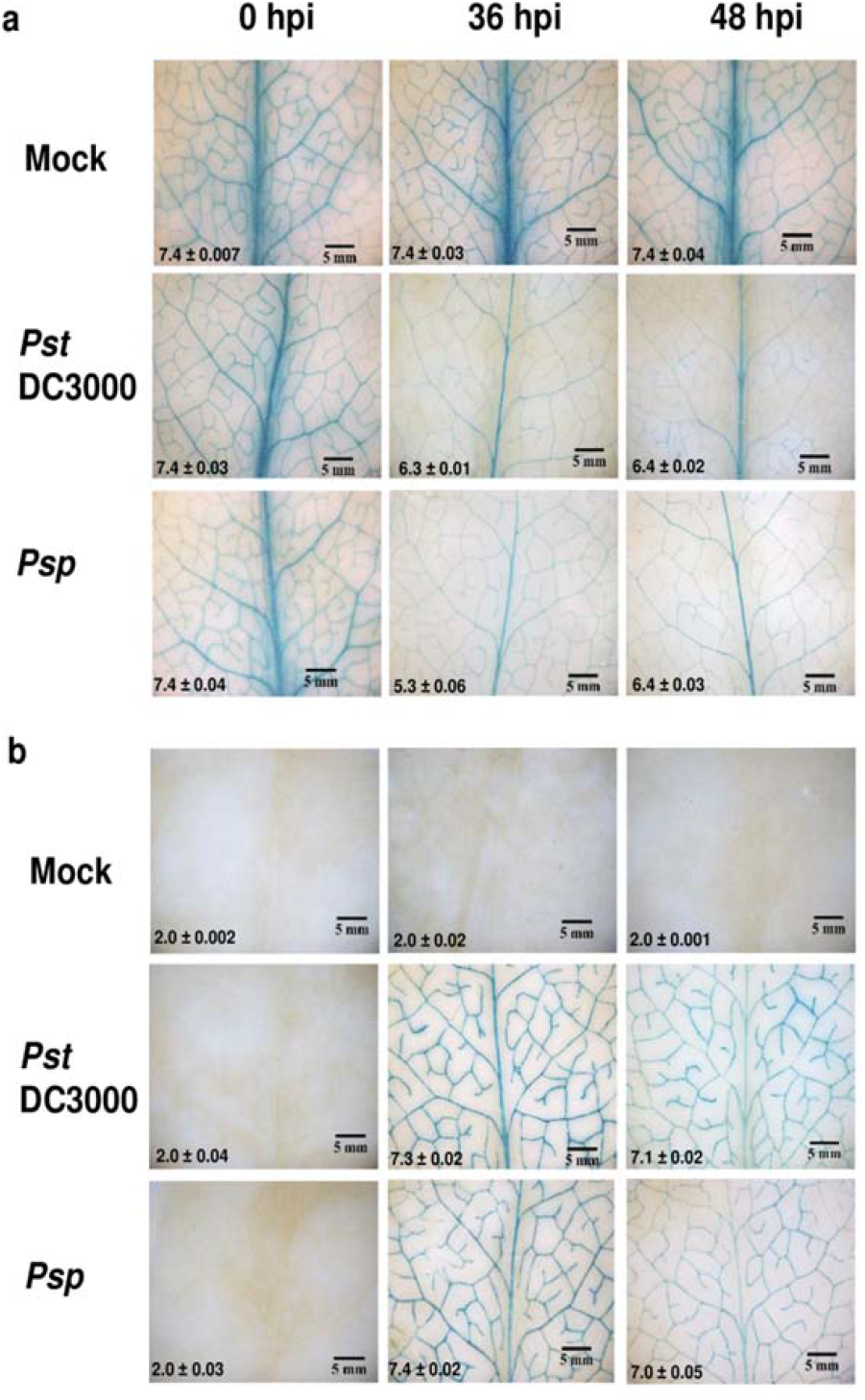
Accumulation of AtSWEET11 and AtSWEET12 transporters after pathogen infection. **a** and **b** The expression of AtSWEET11-GUS and AtSWEET12-GUS translational fusion proteins in the leaf veins was studied by a β-glucuronidase assay. Leaves of 32-day-old Arabidopsis plants expressing stable translational fusion of (**a)** *AtSWEET11:AtSWEET11-GUS* and **(b)** *AtSWEET12:AtSWEET12-GUS* were syringe-inoculated with sterile water (mock), *Pst* DC3000 at 5 × 10^5^ CFU/mL, and *Psp* at 1 × 10^6^ CFU/mL. Samples were collected at 0, 36, and 48 hours post inoculation (hpi). Photographs were taken at 1X magnification after GUS staining, and the stained area was scored by ImageJ software (http://imagej.nih.gov/ij/). The intensity values were calculated and presented at the bottom of the image. The intensity was depicted as log_10_ values obtained from the mean ± SEM of three biological replicates. The experiment was repeated thrice, and consistent results were observed.

In the GUS-based localization study, AtSWEET12 protein was barely detected in the leaves under normal conditions (Fig. 6b). However, the *AtSWEET12* gene was well expressed in the leaves of control wild-type plants (Supplementary Fig. 5). Therefore, we wanted to determine the cellular localization and accumulation status of AtSWEET12 protein in transgenic Arabidopsis plants expressing *p35S:AtSWEET12-eYFP* under normal and pathogen-infected conditions. We found that the AtSWEET12 protein was localized to the plasma membrane in the mock-and pathogen-treated AtSWEET12-YFP overexpression plants (Fig. 7a, Supplementary Fig. 17). Surprisingly, the mock-treated control leaf showed less localization of the AtSWEET12 transporter in the plasma membrane (Fig. 7a). We also observed that many AtSWEET12 proteins were localized to intracellular vesicles in the mock-treated control leaf samples. However, after nonhost and host pathogen infection at 48 hpi, the plasma membrane localization of the AtSWEET12 transporter increased (Fig. 7a). The amount of intracellular localized vesicles retaining AtSWEET12 protein also decreased in pathogen-infected leaves compared to mock-treated leaves (Fig. 7a). Overall, these findings indicate that even after constitutive expression of the *AtSWEET12* gene, plants regulate the abundance and localization of AtSWEET12 at the posttranslational level. During pathogen infection, plants regulate the activity of the AtSWEET12 transporter by increasing the plasma membrane targeting of AtSWEET12 protein to suppress pathogen multiplication. To further confirm that an increase in plasma membrane targeting of AtSWEET12 in an overexpression line restricts bacterial multiplication, we tested the response of an AtSWEET12 overexpression line (expressing *p35S:AtSWEET12-eYFP*) toward the nonhost pathogen *Psp* and host pathogen *Pst* DC3000. The *in planta* bacterial number of *Psp* and *Pst* DC3000 was significantly lower in the AtSWEET12 overexpression line than in the wild type (Fig. 7b,c). *Pst* DC3000-infected AtSWEET12 overexpression plants showed markedly reduced chlorotic symptoms compared to the wild-type plants at 3 dpi (Fig. 7d). These results indicate that *AtSWEET12* overexpression led to plant defense against bacterial pathogens. We further tested the apoplastic sucrose levels in the AtSWEET12 overexpression line and observed an almost 50% reduction in the apoplastic sucrose content compared to the wild type (Fig. 7e). These results indicate that AtSWEET12 controls sucrose availability in the apoplast. Overall, these findings clearly suggest that plasma membrane targeting of AtSWEET12 is required for plant defense against bacterial pathogens by limiting apoplastic sucrose levels.

**Fig. 7.**
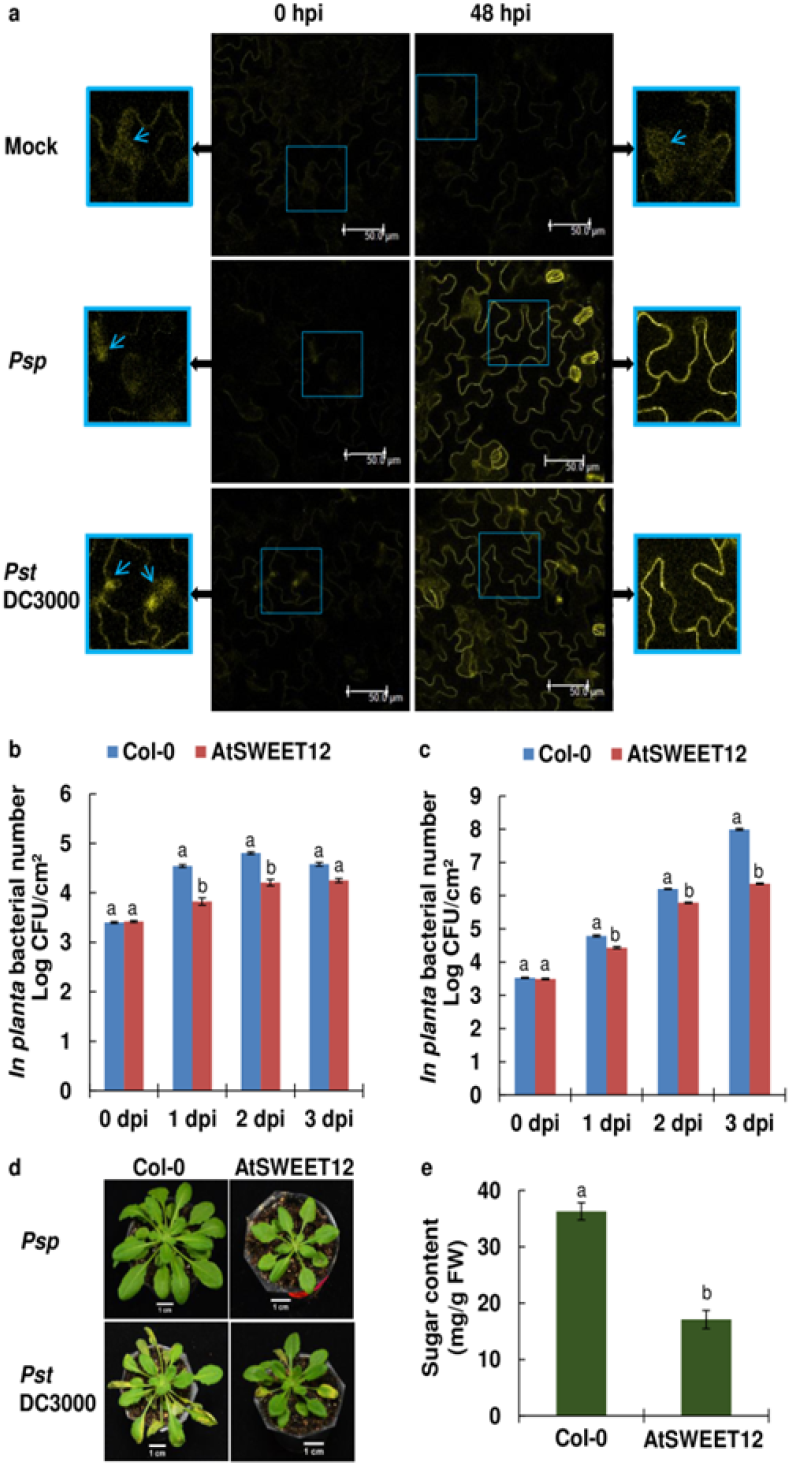
Pathogen exposure triggers plasma membrane targeting of AtSWEET12 protein leading to plant defense. **a** The accumulation of AtSWEET12-YFP fusion proteins was analyzed by confocal microscope after inoculation with sterile water (mock), *Psp* at 1 × 10^6^ CFU/mL and *Pst* DC3000 at 5 × 10^5^ CFU/mL in the leaves of stable transgenic Arabidopsis plants expressing *p35S:AtSWEET12-eYFP*. Samples were collected at 0 and 48 hours post inoculation (hpi). The leaf samples were observed using a Leica TCS SP8 confocal microscope (excitation at 514 nm, fluorescence YFP signal was recorded between 525 and 560 nm). The images were taken using a 63X objective and analyzed by Leica microsystems LAS AF confocal software. The area highlighted by light blue color in the images (middle) was shown in the form of magnified images on left and right side respectively. Light blue color arrow indicates the localization of AtSWEET12-YFP fusion proteins to the intracellular vesicles in the cytoplasm. The bright field images were shown in Supplementary Fig. 16. The experiment was repeated twice, and consistent results were observed. **b** and **c** The leaves of 32-day-old Arabidopsis wild-type (Col-0) and AtSWEET12 overexpression transgenic plants (expressing *p35S:AtSWEET12-eYFP*) were inoculated with **(b)** the nonhost pathogen *Psp* at 1 × 10^6^ CFU/mL and **(c)** the host pathogen *Pst* DC3000 at 5 × 10^5^ CFU/mL. Bacterial multiplication assays were performed, and the bacterial populations were monitored by plating serial dilutions of leaf extracts at 0-, 1-, 2-, and 3-days post inoculation (dpi). *In planta* bacterial number was expressed as log10 values. Data were obtained from the mean of six biological replicates and two technical replicates. Error bars show the standard error of the mean (SEM). **d** The sucrose levels were measured in the apoplast obtained from 32-day-old Arabidopsis wild-type (Col-0) and AtSWEET12 overexpression transgenic plants by a sugar quantification kit. Data were obtained from the mean of five biological replicates (n = 5), and error bars show the SEM. **b, c** and **d** Significant differences (*P* < 0.05) after applying one-way ANOVA and Tukey’s correction are indicated by different letters. The experiment was repeated thrice, and consistent results were observed.

### Heterooligomerization of AtSWEET11 and AtSWEET12 affects sucrose transport

To determine the effect of heterooligomerization of AtSWEET12 with AtSWEET11 on sucrose transport activity, the growth of the sucrose uptake-deficient yeast strain SUSY7/ura3 coexpressing AtSWEET12 and AtSWEET11 was monitored on media containing sucrose as the only carbon source. When AtSWEET12 was coexpressed under a strong promoter (PMA promoter) and AtSWEET11 was expressed from a weaker promoter (ADH promoter) or vice-versa, SUSY7/ura3 growth was dramatically inhibited (Fig. 8a). To confirm the negative affect of heterooligomerization of AtSWEET12 and AtSWEET11 on sucrose transport activity as indicated by the growth assay in SUSY7/ura3 yeast cells, a concentration-dependent sucrose uptake assay was performed in SUSY7/ura3 yeast cells coexpressing AtSWEET12 and AtSWEET11. The analysis of the AtSWEET influx kinetics revealed a 20-fold reduction in V_max_ when AtSWEET12 and AtSWEET11 were coexpressed, provided that AtSWEET12 was expressed under a strong promoter (Fig. 8b, c, e, Supplementary Fig. 18). A five-fold reduction in sucrose influx was observed when AtSWEET12 was expressed from a weaker promoter (Fig. 8d, Supplementary Fig. 18). However, the K_M_ values for AtSWEET11 and AtSWEET12 transporters ranged from 71 mM to 79 mM, but they did not significantly differ from each other (Fig. 8, Supplementary Fig. 18). This inhibition of sucrose transport activity by AtSWEET12 with AtSWEET11 coexpression suggests that the heterooligomerization of AtSWEET12 with AtSWEET11 might be one of the regulatory mechanisms through which AtSWEET11-mediated sucrose efflux could be controlled by AtSWEET12 in the apoplast.

**Fig. 8.**
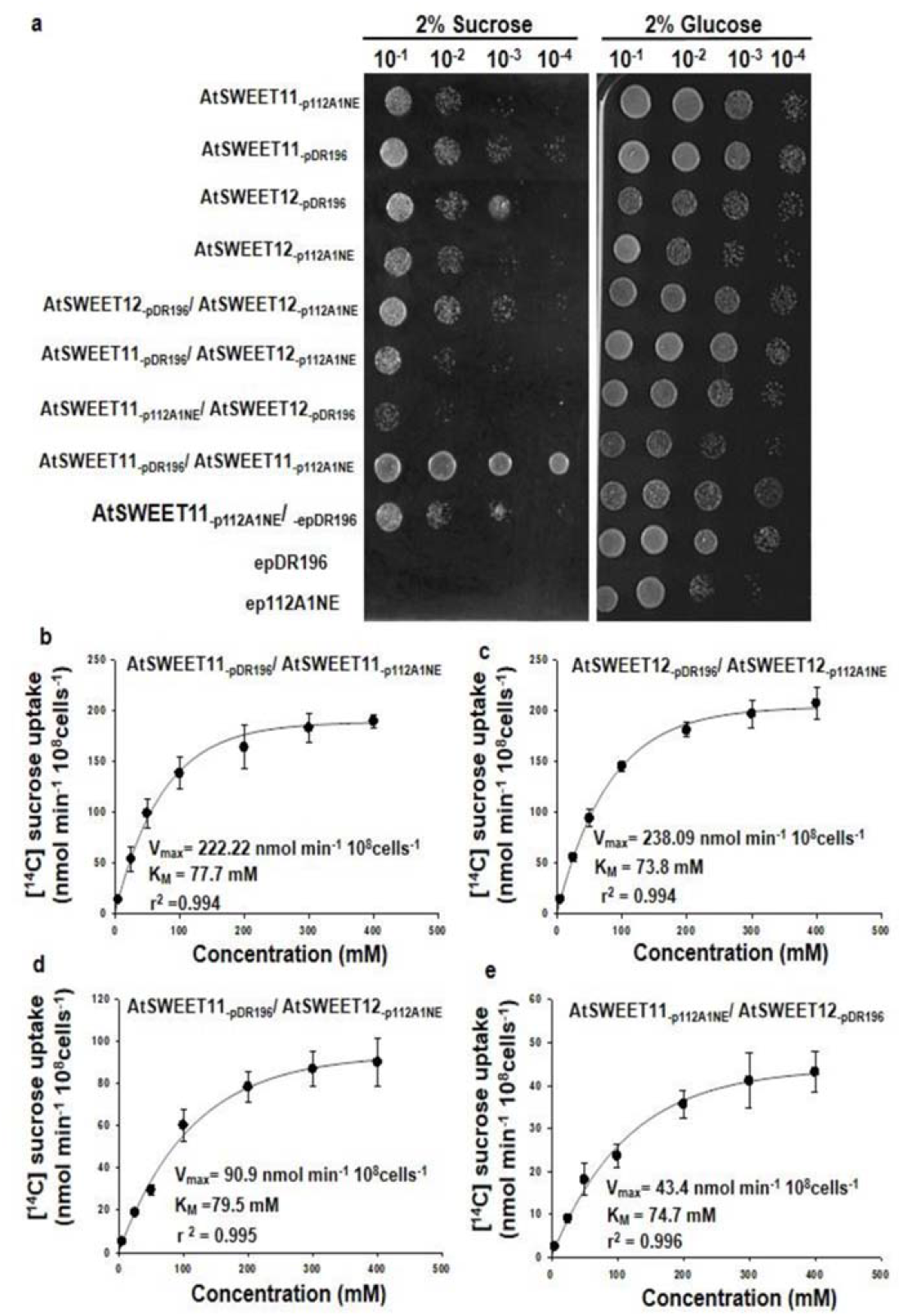
Coexpression of AtSWEET11 and AtSWEET12 transporters in yeast inhibits the sucrose transport activity. **a**, Growth of SUSY/URA3 (sucrose-deficient) yeast transformants expressing *AtSWEET11* and/or *AtSWEET12* either in the vector pDR196 or p112A1NE, and empty vector epDR196 or ep112A1NE on solid media containing 2% sucrose or 2% glucose (control). Images were captured after incubation at 30 °C for 4 days. The experiment was repeated at least three times **b-e**, Concentration-dependent (5mM, 25mM, 50mM, 100mM, 200mM, 300mM and 400mM) [^14^C] sucrose uptake activity was measured in SUSY/URA3 yeast cells coexpressing **(b)** *AtSWEET11* in vector pDR196 or p112A1NE, **(c)** *AtSWEET12* in vector pDR196 or p112A1NE, **(d)** *AtSWEET11* in vector pDR196 and *AtSWEET12* in vector p112A1NE, **(e)** *AtSWEET11* in vector p112A1NE and *AtSWEET12* in vector pDR196. The V_max_, K_M_ and R-squared (**r**^**2**^) values are presented in the graph. The **r**^**2**^ value gives the statistical measure of extent of variation between the two variable in a regression model. Significant differences (P < 0.0001) after applying Student’s t-test, (P < 0.005) after applying one-way ANOVA. Data were obtained from the mean ± standard error (SE) of six replicates (n=6) (see Supplementary Dataset S1 for raw data). The experiment was repeated twice, and consistent results were observed.

## DISCUSSION

The limitation of sugar availability to bacterial pathogens in the apoplast by regulating the sugar transporters involved in sugar efflux and uptake processes has been proposed as a potential plant defense strategy^8,24^. Recently, the role of the sugar uptake transporter AtSTP13—controlling glucose availability to *Pst* DC3000 in the apoplast—was implicated in plant defense^13^. However, the regulation of sugar efflux transporters to limit the sugars to bacterial pathogens has never been reported. Till date, several studies have indicated the modulation of SWEET sugar efflux transporters by pathogens to divert the release of sugars into the apoplast for their nutrition^7,12,19,27,28^. Here, we identified the role of AtSWEET12 in Arabidopsis defense against nonhost and host pathogens by regulating AtSWEET11-mediated sucrose efflux in the apoplast and limiting sucrose availability to bacterial pathogens. These two transporters are known to be involved in the apoplastic loading of mesophyll-synthesized sugars from phloem parenchyma into the phloem apoplast^7^. The effective regulation of these transporters by plants is needed to control the sugar levels in the apoplast, consequently limiting the availability of sugars to pathogens colonizing the apoplast.

Localization experiments using transgenic plants expressing *AtSWEET11*:*AtSWEET11*-*GUS* and *AtSWEET12*:*AtSWEET12*:*GUS* revealed a strong increase in the accumulation of AtSWEET11 protein and low detection of AtSWEET12 protein under normal conditions (Fig. 6). Besides, sugar estimation in the mutant and wild-type plants indicated that the apoplastic sucrose levels decreased only when *AtSWEET11* was mutated, while the sucrose levels remained unaltered when *AtSWEET12* was mutated (Fig. 3a, Supplementary Fig. 11). Thus, AtSWEET11 might be exclusively involved in sucrose transport into the apoplast, while AtSWEET12 is not. This is supported by the induction of *AtSWEET11* expression but not of *AtSWEET12* under high light when endogenous sucrose levels are increased^29^. Thus, AtSWEET11 might be intimately involved in the transport of sucrose to the apoplast during the phloem loading process. Moreover, we predicted a critical role of AtSWEET12 in regulating AtSWEET11-mediated sucrose transport in the apoplast, leading to sucrose limitation during bacterial pathogen infection. The induction of AtSWEET12 and reduction in AtSWEET11 after bacterial pathogen infection, together with the increased pathogen multiplication in the apoplast of *atsweet12* mutants and decreased pathogen multiplication in the apoplast of *atsweet11* mutants, suggest that the control of apoplastic sucrose levels is coordinated by AtSWEET12. This is further supported by the reduced apoplastic sucrose content and stunted phenotype of the AtSWEET12 overexpression line (Fig. 7e, Supplementary Fig. 19). We speculate that due to the overexpression of *AtSWEET12*, the AtSWEET11-mediated sucrose transport in the apoplast is hindered, which affects apoplastic phloem loading and thereby yields the growth defect phenotype of AtSWEET12-overexpressing plants (Supplementary Fig. 19). Besides, AtSWEET12 might be involved in the regulation of AtSWEET11, as both the proteins are co-expressed and interact with each other^6^ (Fig.9; Supplementary Figs. 20–22).

**Fig. 9.**
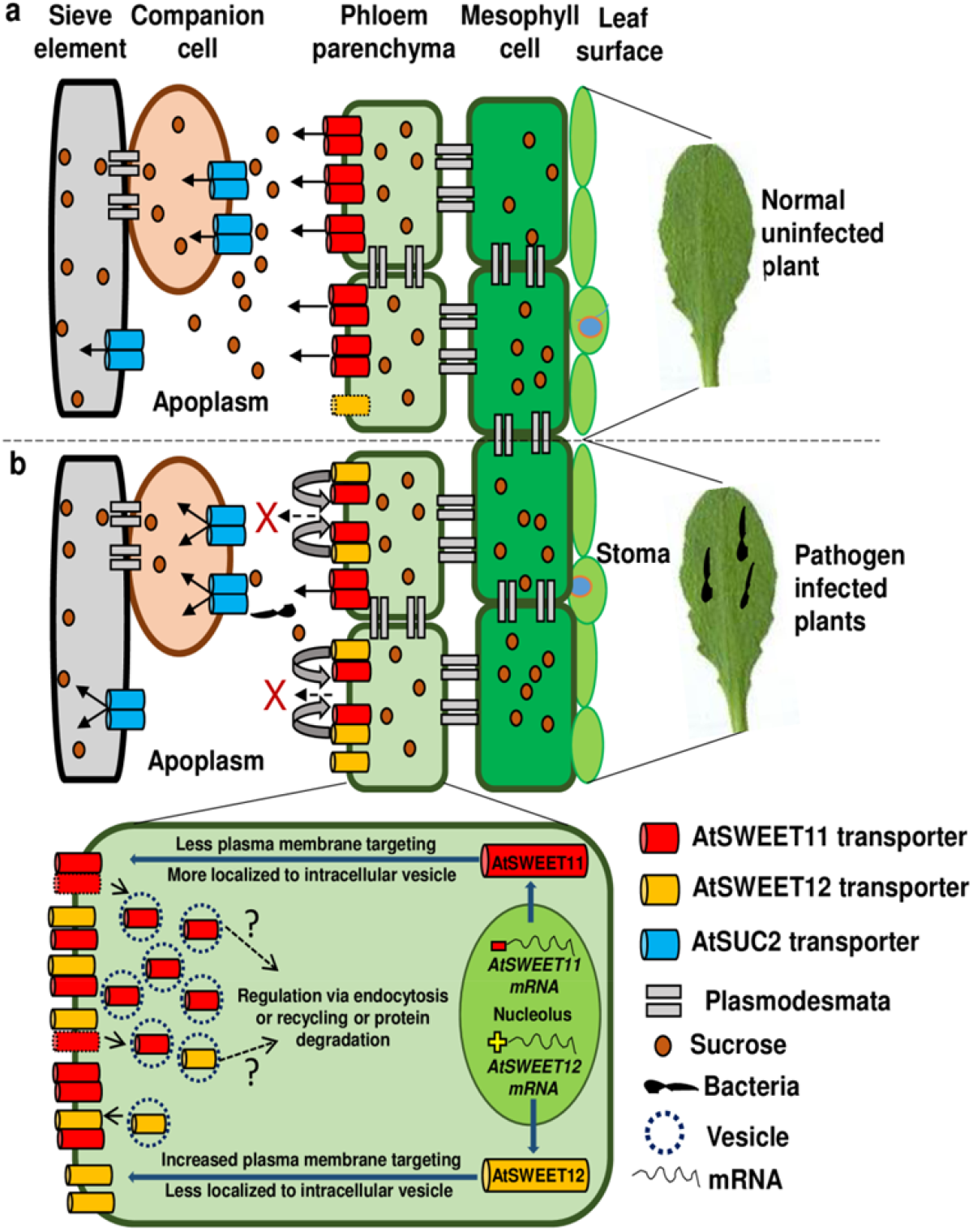
Model depicting the regulation of apoplastic sucrose levels by AtSWEET12 transporter during plant defense. **a** Under normal conditions, AtSWEET11 is actively involved in sucrose efflux into the apoplast during the phloem loading process. **b** After bacterial infection, plant reduces the expression of *AtSWEET11* gene and the abundance of the AtSWEET11 transporter at the plasma membrane. Meanwhile, the plant induces *AtSWEET12* expression and increases the targeting of the AtSWEET12 transporter to the plasma membrane. AtSWEET12 co-interacts and heterooligomerizes with AtSWEET11 that inhibits the transport of sucrose. We speculate that the oligomerization of AtSWEET12 with AtSWEET11 might be one of the regulatory mechanisms through which AtSWEET11-mediated sucrose efflux could be controlled by AtSWEET12, thereby limiting sucrose availability to bacterial pathogens in the apoplast and, thereby leading to the pathogen starvation. Moreover, we propose that plants regulate the localization and abundance of the AtSWEET11 and AtSWEET12 transporters at the plasma membrane by constant endocytosis, recycling, and protein degradation.

Consistent with the *atsweet11;12* double mutant exhibiting reduced susceptibility to the necrotrophic fungus *Colletotrichum higginsianum*^30^ and protist *Plasmodiophora brassicae*^31^, we found that the *atsweet11;12* double mutant exhibited a resistance response towards bacterial pathogens as well (Fig. 2). Moreover, our studies on *atsweet12* mutants and an AtSWEET12 overexpression line confirm the role of AtSWEET12 in plant defense by depriving bacterial pathogens of sugars in the apoplast.The decreased bacterial multiplication and reduced apoplastic sucrose levels in *atsweet11* single and *atsweet11;12* double mutants suggest that AtSWEET11 may contribute to allowing sugar availability to pathogens in the apoplast. The activation of AtSWEET12 and suppression of AtSWEET11 by plant limits sucrose in the apoplast, which restricts pathogen multiplication.

The sugar utilization profiles of *P. syringae* strains indicate that these bacteria prefer sucrose as a nutrient source under *in vitro* conditions^1^ (Supplementary Fig. 12). Accordingly, sucrose limitation might be one of the nonhost resistance strategies employed by plants against nonhost pathogens^2^. Our results validated this hypothesis by demonstrating lower *in vitro* multiplication of bacteria in the apoplast obtained from nonhost pathogen-infected leaves (Fig. 4c). This was further supported by the fact that relieving the sucrose limitation in the apoplast by exogenous addition of sucrose restored *in vitro* bacterial multiplication (Fig. 4d, e). Correspondingly, we found that *in planta* sucrose addition rescued nonhost and host bacterial multiplication in mutants to the wild-type level, which suggests that sucrose is the limiting factor responsible for reduced *in planta* bacterial multiplication in *atsweet11* and *atsweet11;12* mutant plants (Fig. 5). Overall, our study clearly showed that sucrose is crucial for bacterial pathogen multiplication, and that sucrose limitation is one of the defense strategies used by plants to combat nonhost and host bacterial pathogens.

PAMP-mediated activation of STP13 controls apoplastic sugar levels, thereby restricting bacterial pathogen multiplication in the apoplast^13^. Similarly, we found that the plant regulates AtSWEET11 and AtSWEET12 after PAMP perception, as indicated by the strong induction of *AtSWEET12* and down regulation of *AtSWEET11* after treatment with PAMPs and FLG22 and *hrcC*^−^ mutant strains (Supplementary Fig. 5). The plant immune responses are initiated after PAMPs including FLG22 are recognized by pattern-recognition receptors (PRRs) like flagellin-sensitive 2 (AtFLS2) at the plasma membrane. The *in vivo* interaction studies revealed that FLG22 treatment induced the interaction of AtFLS2 with AtSWEET11 or AtSWEET12 which are localized on the plasmamembrane (Supplementary Fig. 23). This indicate that after PAMP perception AtFLS2 may directly modulate the activity of AtSWEET11 and AtSWEET12 proteins. The plant defense strategies may involve the PAMP-mediated regulation of AtSWEET11 and AtSWEET12 transporters that controls the apoplastic sucrose levels to prevent bacterial multiplication.

Besides, basal defense and *R* gene-mediated resistance are largely related, sharing defense pathways, including the salicylic acid (SA)-mediated defense response^32,33^. However, in the *atsweet12* mutant, the *R* gene-mediated resistance against avirulent pathogens remains functional, indicating that the *R* gene-mediated defense pathway is independent of AtSWEET12-mediated plant defense by apoplastic sucrose limitation (Supplementary Fig. 10). Furthermore, SA accumulation and SA-mediated signaling play a crucial role in PTI, contributing to plant defense. The resistance in the *atsweet11;12* mutant was recently attributed to high levels of SA accumulation and the activation of the SA pathway^30^. Consistent with this, we found the upregulation of genes related to SA accumulation and the SA-mediated defense signaling pathway, namely, *enhanced disease susceptibility 1* (*EDS1*) and *pathogenesis-related 1* (*PR1*), in the *atsweet11;12* mutant compared to the wild type (Supplementary Fig. 24). However, SA is not the only factor contributing to the resistance in the *atsweet11;12* mutant. In the *atsweet12* mutant, SA levels were comparable to wild-type levels^30^, and the expression levels of *PR1* and *EDS1* were higher than those in the wild type (Supplementary Fig. 25). Thus, the SA pathway is not affected in *atsweet12* mutants. Through virus-induced gene silencing of *AtSID2*, the SA biosynthesis gene encoding isochorismate synthase in *atsweet12* mutant plants showed more *Pst* DC3000 multiplication and higher sucrose levels than in *atsweet12* mutant plants (Supplementary Figs. 25, 26). Given these findings, SA-mediated signaling could be one of the upstream components but not directly involved in regulating sucrose availability in the apoplast in response to pathogen infection.

Moreover, the regulation of transporters may occur at the posttranslational level^26^. YFP-based localization in the AtSWEET12 overexpression line revealed that under normal conditions, AtSWEET12 is retained inside the intracellular vesicle rather than at the plasma membrane, and that pathogen infection triggers more plasma membrane targeting of AtSWEET12 (Fig. 7a). Accordingly, we surmise that Arabidopsis might regulates the localization and abundance of AtSWEET12 at the plasma membrane by constant endocytosis, recycling, and protein degradation. Future studies are needed to explore the mechanisms involved in the regulating the abundance of AtSWEET12 proteins. Further, oligomerization has been identified to be necessary for the function of AtSWEETs, and AtSWEET11 homooligomerizes to form a functional pore that allows the transport of sucrose^34^. Besides, the oligomerization of functional AtSWEET1 with defective AtSWEET1 inhibits glucose transport activity^34,35^. In another study, the oligomerization of functional OsSWEET11 with the mutated form of OsSWEET11 inhibited the sugar transport activity and restricted *Rhizoctonia solani* infection in rice^21^. Biochemical studies have indicated that the heterooligomerization of the sucrose transporters SUT1 and SUT2 ceases sucrose transport^36^. Moreover, AtSWEET12 and AtSWEET11 co-interact with each other (Fig. 9 and Supplementary Figs. 20–22) and have been reported to form a heterooligomer ^6,34^. Our data provide evidence that the coexpression of AtSWEET12 with AtSWEET11 in yeast cells inhibited sucrose uptake activity (Fig. 8). This might be one of the predictable mechanisms for AtSWEET12-mediated regulation of AtSWEET11 through which sucrose levels can be controlled in the apoplast during bacterial infection. We suspect that the heterooligomerization of AtSWEET12 with AtSWEET11 may hinder the transport activity because structural incompatibility prevents the formation of a functional pore for sucrose transport.

Various studies lead us to state that the plant defense strategy of apoplastic sucrose limitation to bacterial pathogens might involve the highly coordinated regulation of AtSWEET11 and AtSWEET12, and the sugar uptake transporters AtSUC2 to restrict bacterial pathogen multiplication^6,8,13^. On the basis of our findings, we propose that under normal conditions, AtSWEET11 is involved in sucrose efflux into the apoplast. During bacterial infection, plant induces *AtSWEET12* and reduces *AtSWEET11* expression after PAMP perception, and promotes the plasma membrane targeting of AtSWEET12 protein. The AtSWEET12 heteroligomerize with AtSWEET11 which inhibit the sucrose transport, thereby limiting sucrose availability to the pathogen in the apoplast and restricting its multiplication (Fig. 9). Besides, phosphorylation and ubiquitination might be involved in the activation and deactivation of these transporters^13,37,38^. The interaction of AtFLS2 with AtSWEET11 or AtSWEET12 could be directly involved in mediating the post translational regulation of these transporters. In the road ahead, more focused studies exploring the mechanisms involved in the regulation of AtSWEET11 and AtSWEET12 will provide clearer insights into this aspect as a basis of plant defense. Taken together, our findings clearly showed a plant defense strategy against bacterial pathogens involving the PAMP-mediated regulation of AtSWEET11 and AtSWEET12. Our findings highlight a role of AtSWEET12 in controlling apoplastic sucrose levels, thereby restricting bacterial multiplication in the apoplast. Moreover, along with other active plant defense mechanisms, the regulation of apoplastic sugar availability to pathogens at the site of infection by manipulating such sugar transporters can be an effective strategy for crop protection.

## MATERIALS AND METHODS

### Plant material and growth conditions

*Arabidopsis thaliana* wild-type Columbia-0 (Col-0); T-DNA mutants of all 17 *AtSWEET* genes (*atsweet1* to *atsweet17*) (described in Supplementary Table 2; more details regarding genotyping and null mutation analysis are presented in Supplementary Materials and Methods, Supplementary Figs. 27 and 28, and Supplementary Tables 2 and 3); *atsweet11;12, atsweet11;15, atsweet12;15, atsweet11;12;15* mutant lines^17^, and complementation lines of AtSWEET11 (c-11/11;12) and AtSWEET12 (c-12/11;12) in the *atsweet11;12* double mutant background^6^; *pAtSWEET11:AtSWEET11-GUS* and *pAtSWEEt12:AtSWEET12-GUS* lines^6^; and *p35S:AtSWEET12-eYFP* (overexpression line) were obtained from the Arabidopsis Biological Resource Centre (ABRC) (https://abrc.osu.edu/). For seedling germination and growth, Arabidopsis seeds were sown in a soil mixture of 3:1 vol/vol agropeat (Prakruthi Agro Tech, Bangalore, India) and vermiculite (Keltech Energies Ltd., Bangalore, India) and then cold treated for 3 days at 4 °C in the dark. Arabidopsis plants were grown in a growth chamber (PGR15; Conviron, Winnipeg, Canada) under 8 h light (light intensity, 200 μE m^−2^ s^−1^)/16 h dark at 20 °C and 75% relative humidity. Plants were irrigated alternately with water and 1/2X Hoagland nutrient solution (Cat# TS1094; HiMedia Laboratories, Mumbai, India) every day.

### Bacterial pathogens and inoculum preparation

The bacterial pathogens used in this study were as follows: a host pathogen of Arabidopsis, *Pst* DC3000; nonhost pathogens of Arabidopsis, *Psp* and *Pta*; T3SS mutants (*hrcC*^−^) of all these three bacterial pathogens; and avirulent strains of *Pst* DC3000, i.e., avrRpt2, avrRps4, and avrRpm1. All the bacterial strains were grown at 28 °C with continuous shaking at 150 rpm in King’s B (KB) medium (liquid) (Cat# M1544; HiMedia Laboratories) containing appropriate antibiotics. Rifampicin at 50 μg/mL was added to the medium for growing *Pst* DC3000 and *Psp*. Bacterial cultures were grown overnight (12 h) to obtain an optical density of 0.4 at 600 nm (OD_600_ = 0.4). Bacterial cells were collected by centrifugation at 4,270 *×g* for 10 min, washed thrice in sterile water, and re-suspended in sterile water at desired concentrations. The concentrations used for the inoculation of the leaves (32-day-old plants) were 5 × 10^5^ colony-forming units (CFU)/mL for *Pst* DC3000, 1 × 10^6^ CFU/mL for *Psp*, 3 × 10^5^ CFU/mL for *Pta*, and 1 × 10^6^ CFU/mL for the *hrcC*^−^ T3SS mutants of *Pst* DC3000 and *Pta*.

### Plant inoculation, *in planta* bacterial multiplication assay, and disease index analysis

Thirty-two–day-old plants were used for inoculation experiments. The bacterial suspension was syringe-infiltrated on the abaxial surface of fully expanded leaves using a needleless syringe. Each plant (all leaves) was inoculated with 5 mL of bacterial suspension. Sterile water-infiltrated plants were used as mock plants. For the estimation of bacterial multiplication after *in planta* sucrose addition, the bacterial pathogen was co-infiltrated with sucrose at a low concentration of 0.2% (2 mg/mL) and a high concentration of 1% (10 mg/mL). The inoculated plants were maintained in a growth chamber at 20 °C. Leaf samples were collected at 0, 1, 2, and 3 dpi for bacterial multiplication analysis. *In planta* bacterial multiplication levels were quantified by homogenizing leaf discs (0.5 cm^2^ each) in sterile water, after which the appropriate dilution was plated on KB agar medium. Bacterial numbers were counted as CFUs per square centimeter of leaf area and expressed as log_10_ values^39^. Experiments were carried out with at least six plants per treatment as biological replicates, and two leaves from each plant were taken as technical replicates. Bacterial multiplication numbers were calculated by using the following formula:

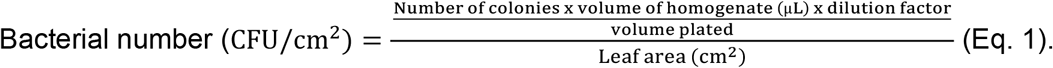

Log_10_ values are presented in the graphs. The disease index after host pathogen infection was established by scoring the chlorotic and necrotic disease symptoms in plants at 3 dpi. Five different plants were used as biological replicates for scoring the disease symptoms. Photographs were taken at 3 dpi.

### Green and yellow fluorescence detection by confocal microscopy

The imaging of *in planta* bacterial populations of *Pst* DC3000 and *Pta* carrying the pDSK-GFPuv plasmid^39^ was done at the cellular level using a confocal microscope (Leica TCS SP8 AOBS system; Leica Microsystems, Wetzlar, Germany). A small piece of leaf from 32-day-old Arabidopsis wild-type and mutant plants inoculated with GFPuv-labeled bacteria was kept on a glass slide. The leaf tissue was submerged with a droplet of deionized water and covered with a glass cover slip. The images were taken with a water immersion objective of 63X using excitation at 488 nm and emission between 500 and 600 nm. Similarly, localization of AtSWEET12-YFP in Arabidopsis leaves was performed using the same instrument after excitation at 514 nm, and the fluorescence YFP signal was recorded between 525 and 560 nm.

### GUS histochemical staining and microscopic analysis

Histochemical GUS staining was performed by following the standard protocol as explained by Jefferson et al.^40^(1987). Leaf samples were harvested after pathogen inoculation and mock treatment at 0, 36, and 48 hpi. The leaf tissues were pre-incubated in 90% (v/v) ice-cold acetone for 10 min and then washed thrice with 100 mM phosphate buffer solution (pH 7.2) for 5 min. Leaf tissues were then incubated at 37 °C for 8 h in a GUS staining solution containing 1 mM of 5-bromo-4-chloro-3-indolyl-β-glucuronic acid, cyclohexylammonium salt monohydrate (X-Gluc) (Cat# B-7300; Biosynth AG, Staad, Switzerland), 100 mM sodium phosphate (pH 7.0), 1 mM potassium ferrocyanide, 1 mM potassium ferricyanide, and 0.1% (v/v) Triton X-100 (Cat# MB031; HiMedia Laboratories). Gradient alcohol treatment was performed for de-staining the leaf tissue samples. The leaf samples were first immersed in 90% alcohol for 4 h, followed by 70% alcohol for 12 h, 50% alcohol for 24 h, and 30% alcohol for 48 h. The GUS-stained samples were observed under a Strereozoom AZ100 Microscope (Nikon Instruments Inc., Melville, NY, USA) mounted with a digital camera (Nikon Digital Sight DS-Rs1; Nikon Instruments Inc.) for capturing the images. Fiji software (http://fiji.sc) was used for image analysis. ImageJ software (http://imagej.nih.gov/ij/) was used for measuring the intensity of the images.

### Apoplastic fluid extraction

Apoplastic fluid was extracted from control and pathogen-infected Arabidopsis plants according to the standard protocol detailed by O’Leary et al.^41^(2014), with minor modifications. Arabidopsis leaves were detached and rinsed in distilled water to remove surface contamination. Leaf weight was measured before infiltration, after which leaves were immersed in a beaker containing 250 mL sterile MilliQ water.

Vacuum was applied to infiltrate sterile water into the apoplastic space till complete saturation. The leaves were blotted dry, weighed, and placed in 4-inch-wide Parafilm strips and gently rolled up. The rolled-up leaves were inserted into a 20 mL syringe, and this setup was further placed into a 50 mL centrifuge tube for spinning. Apoplastic fluid extracted by centrifugation at 1,000 *×g* for 30 min at 4 °C was transferred into 1.5 mL tubes and centrifuged at 15,000 *×g* for 5 min to remove particulate materials. The apoplastic fluid was then filter-sterilized through 0.22 μm filters to remove any microbial contamination. No cytoplasmic contamination was detected in the apoplastic samples (detailed information is presented in Supplementary Materials and Methods and Supplementary Fig. 29). Samples were then used for further analysis or stored at −80 °C.

### GC-MS metabolite analysis

Metabolite profiling of apoplastic fluid obtained from infected and control plants was done by following the standard protocol of Lisec et al.^42^ (2006) with minor changes. Apoplastic fluid samples (500 μL) were transferred into 2 mL tubes, and the samples were lyophilized in a freeze drier overnight. A calculated amount of distilled water was added to reconstitute the apoplastic sample. Next, 200 μL apoplastic sample was transferred to 1.5 mL tubes, and 700 μL methanol with 15 μg/mL adonitol (Cat# 02240; Sigma-Aldrich, St. Louis, USA) as the internal standard was added, followed by incubation at 70 °C for 10 min. Then, samples were centrifuged at 11,000 *×g* for 10 min, and 700 μL of supernatant was transferred to fresh 1.5 mL tubes. Ice-cold chloroform (375 μL) and molecular-grade water (500 μL) were added to the supernatant, and the mixture was vortexed and centrifuged at 2,200 *×g* for 15 min. Then, 150 μL of the upper aqueous layer was transferred to 4 mL glass vials and dried completely by vacuum centrifugation at room temperature. The methoximation of dried samples was done by adding 40 μL pyridine (Cat# 270407; Sigma-Aldrich) with 20 mg/mL methoxyamine hydrochloride (Cat# 89803; Sigma-Aldrich) and incubating at 70 °C for 2 h with vigorous shaking. The sample was then derivatized by adding 60 μL *N*-methyl-*N*-(trimethylsilyl)trifluoro acetamide (MSTFA) (Cat# 69479; Sigma-Aldrich) and incubating for 30 min at 37 °C. Two microliters of sample was used for analysis by an autosampler-autoinjector (AOC-20si) coupled with a gas chromatograph-mass spectrometer (Shimadzu QP2010 Ultra, Kyoto, Japan).

Analyses of chromatograms and mass spectra were performed by GCMS solution software (Shimadzu). The peaks were identified as metabolites using spectral libraries, i.e., NIST8 and WILEY8. The peak area for each metabolite was normalized using adonitol as the internal control.

### Estimation of sugar content

Five hundred microliters of apoplastic fluid was transferred into 2 mL tubes, and the samples were lyophilized in a freeze drier overnight. The apoplastic fluid sample was reconstituted with distilled water. Next, 200 μL of concentrated apoplastic sample was transferred to 1.5 mL tubes. Soluble sugars were extracted by adding 500 μL of 80% ethanol at 80 °C for 10 min. Samples were centrifuged at 11,000 *×g* for 10 min, and 300 μL of supernatant was transferred to fresh 1.5 mL tubes. Sucrose, D-fructose, and D-glucose concentrations were determined from the apoplastic samples by using a Sucrose/D-Glucose/D-Fructose Megazyme Kit following the manufacturer’s instructions (K-SUFRG; Megazyme International, Ireland Ltd., Wicklow, Ireland). Sugars were quantified by reading the samples at 340 nm using a spectrophotometer.

### *In vitro* bacterial quantification

*Pst* DC3000, *Psp*, and *Pta* were grown in KB medium overnight (12 h) till the initial optical density reached OD_600_ = 0.4. Bacterial cells were harvested by centrifugation at 5,488 *×g* for 10 min. Pellets containing bacterial cells were washed twice with sterile water and re-suspended in sterile water. Bacterial suspensions were inoculated in MM (pH 7.4) supplemented with 5% (v/v) apoplastic extracts from mutants and wild-type plants individually, such that the OD_600_ at the 0 h time point was maintained at 0.1. The samples were then grown at 28 °C with continuous shaking at 150 rpm. Bacterial quantification was performed at 0, 8, 10, and 12 h by measuring the OD_600_ for each sample.

### RT-qPCR analysis

The bacterial cells were harvested, and the total RNA was extracted using TriZol reagent (Cat# 15596018, Invitrogen, Carlsbad, CA, USA) as per the manufacturer’s protocol. RNA quantification was done using a NanoDrop spectrophotometer (ND-1000; Thermo Fisher, Waltham, MA, USA). RNA (5 μg) was treated with DNase. First-strand cDNA synthesis was done using DNase-treated RNA in a reaction volume of 50 μL by a Verso cDNA synthesis kit (Cat# AB1453A, Thermo Scientific) following the manufacturer’s protocol. The gene-specific primers, including the primers for the reference gene *16S rRNA* (Psp: PSPPH_0689; Pst DC3000: PSPTOr01), were designed using Primer 3 software (http://bioinfo.ut.ee/primer3-0.4.0/). Details of the primers used in the study are listed in Supplementary Table S3^43^. For real-time quantitative PCR (RT-qPCR), a 10 μL final volume was prepared by adding 1 μL of five-fold diluted cDNA, gene-specific primers at 750 nM each, and HotStart-IT SYBR Green qPCR Master Mix (Cat# 600882, Agilent Technologies, Santa Clara, CA, USA) as per the manufacturer’s protocol. RT-qPCR was performed according to the manufacturer’s instructions on an ABI 7900HT PCR system (Applied Biosystems, Foster City, CA, USA). The cycle threshold (Ct) values for *16S rRNA* expression were used to normalize the expression values of target genes in each sample. The relative expression values for each sample were determined over their respective control using the comparative 2^−ΔΔCt^ method ^44^. Three independent biological replicates were used for all RT-qPCR analyses.

### Yeast growth assay

Cells from single colonies of the transformed yeast strain SUSY7/ura3 ^45^ were grown overnight at 30 °C. A portion of this was re-inoculated into fresh synthetic deficient (SD) medium with 2% glucose until the cell densities reached an OD_600_ of 0.8–1.0. Cells were then harvested by centrifugation at 4,270 *×g* for 10 min, washed twice, and then diluted with SD medium (without any carbon source) to an OD_600_ of 0.2. Next, the appropriate serial dilutions of all desired yeast cells were prepared (10^−1^, 10^−2^, 10^−3^, 10^−4^), and from every dilution, 5 μL was spotted on the plates containing SD medium supplemented with 2% glucose or 2% sucrose. Plates were kept for incubation at 30 °C for 4 days and then scanned and photographed (detailed information about constructs is presented in Supplementary Materials and Methods).

### Radiotracer sucrose uptake assay in yeast

Cultures of transformed SUSY7/ura3 yeast cells were grown overnight at 30 °C. A portion of this culture was diluted to an OD_600_ of 0.2 with fresh SD medium supplemented with 2% glucose and kept for 4 h incubation at 30 °C until the cell densities reached an OD_600_ of 0.5. Cells were collected by centrifugation, washed twice, and then re-suspended into 50 mM sodium phosphate buffer (pH 5.5) to an OD_600_ of 10. The concentration-dependent sucrose uptake assay was performed as described by Sauer and Stadler^46^(1993) and Ho et al.^47^(2019). The uptake buffers were prepared with desired sucrose concentrations in 50 mM sodium phosphate buffer with equimolar ratio of [^14^C]-radiolabeled sucrose. The uptake assay was initiated by adding equal volume of transformed yeast cells into 150 μL of uptake buffers and then incubated at 30 °C for 10 min. Yeast cells were collected by vacuum filtration on MCE membrane filter paper (pore size, 0.45 μm) and then washed thrice with 5 mL of ice-cold 50 mM sodium phosphate buffer. The filter paper containing cells was kept in 5 mL of counting fluor (Sigma) in scintillation vials. Radioactivity was quantified by a liquid scintillation counter (Perkin Elmer Inc., USA), and kinetic analysis for nonlinear regression was performed using SigmaPlot (version 14; Systat Software Inc., Chicago, IL, USA).

### Statistical analysis

The error bars presented in the figures represent the standard error of the mean (SEM) of all the replicates used in the study. The figure legends state the total number of biological replicates used in every experiment. For statistical analysis of data represented in the figures, two-way and one-way analyses of variance (ANOVA), along with Tukey’s post-test comparison (*P* < 0.05), were performed using SigmaPlot (version 14.0). Student’s *t-*test at *P* < 0.05 was performed to determine significant differences from the wild type. The raw data containing the details of the total number of biological replicates and SEM values are presented in Supplementary File 1.

## Data availability

All data related to this study are available within the manuscript and its supplementary files or are available from the corresponding author upon reasonable request. The raw data containing the details for figures and supplementary figures are provided in Supplementary File 1.

## Accession Numbers

For this article, the sequence data can be retrieved from the Arabidopsis Genome Initiative or GenBank/EMBL databases under the following accession numbers: *SWEET1*: *AT1G21460, SWEET2*: *AT3G14770, SWEET3*: *AT5G53190, SWEET4*: *AT3G28007, SWEET5*: *AT5G62850, SWEET6*: *AT1G66770, SWEET7*: *AT4G10850, SWEET8*: *AT5G40260, SWEET9*: *AT2G39060, SWEET10*: *AT5G50790, SWEET11*: *AT3G48740, SWEET12*: *AT5G23660, SWEET13*: *AT5G50800, SWEET14 AT4G25010, SWEET15*: *AT5G13170, SWEET16*: *AT3G16690, SWEET17*: *AT4G15920, SID2*: *AT1G74710, PR1*: *AT2G14610, PR5*: *AT1G75040, EDS1*: *AT3G48090, scrY*: *PSPPH5187, PSPTO0890*.

## Acknowledgements

We thank Dr. W.B. Frommer and the Arabidopsis Biological Resource Center for providing seeds for transgenic plants. We acknowledge Drs. John Ward and Divya Chandran and for sharing yeast strains. We thank Ms. Nishtha Rawat and Ms. Anjali for technical help during experiments. We thank Mr. Rahim Tarafdar and Sundar Solanki for providing technical help at the laboratory and Dr. Aashish Ranjan, Dr. Senjuti Sinharoy, Dr. Mahesh Patil, Dr. Bendangchuchang Longchar, Dr. Piyush Priya, and Aanchal Choudhary for critical reading of the manuscript. We acknowledge the DBT-eLibrary Consortium (DeLCON) and NIPGR library for providing access to e-resources and the NIPGR Plant Growth Facility for plant growth support/space. The project at MS-K’s laboratory was funded by NIPGR. UF acknowledges the DBT-SRF fellowship (DBT/2013/NIPGR/68) and NIPGR-SRF fellowship.

## Author contributions

MS-K conceived the idea. MS-K and UF designed the study. UF executed the experiments, analyzed the data, and contributed to drafting the manuscript. MS-K and UF edited the manuscript.

## ADDITIONAL INFORMATION

